# Establishment of a cochlear lymphatic vascular network and immune surveillance system following cochlear implantation

**DOI:** 10.64898/2026.10.22.760829

**Authors:** Muhammad T. Rahman, Ibrahim Razu, Shakila M Fatima, Brian J Mostaert, Bryce Hunger, Md. F. Hossain, Diana Correa, Jennifer O’Malley, Nashwaan A. Khan, Zhenshen Zhang, Osama Tarabichi, Ya Lang Enke, Robert D. Gay, Jacob Oleson, Noah S. Butler, Young-Kwon Hong, Alicia Quesnel, Alexander D. Claussen, Marlan R. Hansen

**Affiliations:** Department of Otolaryngology-Head and Neck Surgery, The University of Iowa, Iowa City, IA-52242, USA; Mass Eye and Ear 243 Charles Street Boston, MA 02114; Department of Biology, The University of Iowa, Iowa City, IA 52242, USA; Cochlear Limited, Sydney, Australia; Department of Biostatistics, The University of Iowa, Iowa City, IA-55542, USA; Department of Microbiology and Immunology, The University of Iowa, Iowa City, IA-52242, USA; Department of Surgery, Beth Israel Deaconess Medical Center, Harvard Medical School, Boston, MA

**Keywords:** Lymphatics, immune surveillance system, cochlear implant, foreign body response

## Abstract

Hearing loss is a highly prevalent sensory abnormality often addressed with cochlear implants (CI) that provide auditory rehabilitation for deaf patients. However, a universal inflammatory foreign body response (FBR) to the implanted materials negatively impacts CI outcomes. The mechanistic basis for immune cell infiltration and propagation of the FBR directed against CIs is not defined. Using mouse CI models and quantitative, highly tractable reporters for lymphatics, blood vessels, and immune cells, we identify inner ear lymphatic vessels and changes to the immune surveillance system following cochlear implantation. Histopathological analysis of implanted temporal bones from human subjects shows that both blood vessels and lymphatics are evident within the scala tympani of the implanted cochlea. Our data reveal a rich lymphatic vascular network in the inner ear, closely associated with the perilymphatic and endolymphatic spaces. Lymphatics proliferate within the perilymphatic space of the implanted cochleae and appear to traffic immune cells. Together, our data provide direct evidence that concurrent proliferation of blood vessels and recruitment of blood-borne immune cells comprise an ‘immune surveillance system’ in the implanted cochleae that promotes the FBR. Our findings also highlight potential opportunities to limit adverse FBR outcomes following cochlear implantation.

## 1. Introduction

Hearing loss is a highly prevalent sensory abnormality with nearly 1.57 billion people affected globally in 2019, which is projected to affect ∼2.45 billion people by 2050.(1) In patients with profound to severe sensorineural hearing loss, cochlear implants (CI) provide auditory rehabilitation, improve cognitive function, and quality of life,(2–4) as well as reduce mental illness(5, 6) and health care and educational costs.(4, 7) An inflammatory foreign body response (FBR), universally observed following CI surgery, has been linked with poorer performance and failure of CI, loss of residual acoustic hearing, post-implant degeneration of hair cells, spiral ganglion neurons (SGNs), and increased electrode impedance in humans and animal models.(8) Understanding the immune response to cochlear implantation(8) requires a detailed characterization of the cochlear ‘immune surveillance system’.(9, 10) ‘Immune surveillance’ requires two-way trafficking of immune cells, proteins, and immune complexes through blood and lymphatic vessels.(11, 12) Under basal conditions, fluid transudates from the blood vessels into the interstitial space, generating a directional fluid flow and prompting passive immune cell transmigration through discontinuous junctions of lymphatic capillaries.(10, 13) Tissue resident antigen-presenting cells (APCs) phagocytose antigens(14) migrate to the draining lymph node(s) through afferent lymphatic vessels and present antigens to CD4+ T lymphocytes aided by the antigen-presenting molecule, major compatibility complex (MHC) II. Dendritic cells play a critical role as APCs, activating CD4+ T cells(10) that subsequently exit the lymph node and reach the systemic blood circulation via the efferent lymphatic vessels, infiltrate the tissue via the blood circulation, and orchestrate the activity of other immune cells.(9) In settings of both infection and sterile tissue injury, fluid flow and trafficking of immune cells to and from damaged tissue are accelerated,(10) contributing to chronic inflammation.(15)

Major components of a putative ‘immune surveillance system’ of the cochlea that include the blood supply(16) and immune cells(17) have been extensively characterized, yet little is known about whether a cochlear lymphatic vascular system exists. In an experiment using a leukocyte tracer a link between the cochlea and the lymphatic system has been suggested.(18) The presence of molecular markers of lymphatics in the mouse cochlea,(19) evidence for lymphatics in pathological human cochleae,(20) and other indirect evidence suggest that a lymphatic network exists within the cochlea, draining the organ.(17) However, characterization of the lymphatic vasculature network in undamaged cochleae has not been directly demonstrated. Likewise, the response of cochlear lymphatics and their relationship to the inflammatory foreign body response (FBR) following implantation remains unknown. In the current study, we investigated the presence and function of the cochlear lymphatic system and its role in the inflammatory FBR following cochlear implantation. We describe dynamic alteration of lymphatics, blood vessels, as well as the redistribution and infiltration of mononuclear antigen-presenting cells (e.g., dendritic cells), non-macrophage immune cells, and T lymphocytes that required angio- and lymphangiogenesis. Collectively, our data reveal a rich lymphatic vascular network in the inner ear that is activated during the FBR to cochlear implantation.

## 2. Results

We first investigated the spatial organization of different components of the immune surveillance system (lymphatics, blood vessels, and immune cells) within the mouse cochlea using transgenic reporter mice engineered to report lymphatics, blood vessels, and immune cells. Then, we investigated how these components respond to cochlear implantation in mouse and human cochleae. The results of these experiments are summarized.

### 2.1. Microscopic evidence for mouse cochlear lymphatics

#### 2.1.1. Cochlear lymphatic markers in cryosections

Podoplanin/PDPN, Lyve1, VEGFR3, and Prox1 are molecular markers that have been commonly used to define lymphatic vasculature in various organs, including the central nervous system.(21) To identify cochlear lymphatics using confocal imaging, we first optimized the immunolabeling protocol using small intestine villi using Prox1-GFP and Prox1-tdTomato lymphatic reporter mice (supplementary materials, Supplementary Figure 1). To distinguish lymphatic vessels from blood vessels, we generated a lymphatic-blood vessel dual reporter mouse (Prox1-tdTomato: PECAM1-GFP) (supplementary materials, Supplementary Figure 1). Further validation of lymphatic markers in the cochlea is described in the supplementary material (Supplementary Figure 2). Our findings show that while Prox1 is considered to be a sensitive marker for lymphatic vessels, it lacks the specificity within the cochlea as it is expressed in other cells (e.g., SGNs). Therefore, while we employed Prox1-GFP and Prox1-tdTomato transgenic reporter mice for this study, we recognized Prox1+ cells as lymphatics only when they simultaneously expressed additional markers of lymphatics (e.g., PDPN, Lyve1, or VEGFR3).

To identify lymphatic vessels in the cochlea, we labeled cryosections with an antibody against PDPN. This revealed distinct PDPN+ vascular structures within the spiral ganglion, stria vascularis, and along the margins of ST and scala vestibuli (Figure 1, Supplementary Video 1, and Supplementary Video 2). Sections from the cochleae from lymphatic reporter (Prox1-GFP) mice labeled with anti-PDPN, anti-VEGFR3, or anti-Lyve1 antibodies revealed that Prox1 co-localized with PDPN, VEGFR3, and Lyve1 in regions of the cochlea adjacent to the perilymph spaces: ST and RWM, indicating their identity as lymphatic vessels (Figure 1). In lymphatic-blood vessel dual reporter (Prox1-tdTomato: PECAM1-GFP) mice, Prox1+ cells in the cochlea were distinct from PECAM1+ blood vessels (Figure 1 and Supplementary Video 3). Further, in sections from PECAM1-GFP mice labeled with anti-PDPN antibody, PDPN+ lymphatic vessels were found to be distinct yet adjacent to PECAM+ blood vessels throughout the cochlea. (Figure 1 and Supplementary Figure 3). Spatially, the lymphatic vascular network was widely distributed throughout the cochlea, with a high density of lymphatic vessels close to the vascular planes (outer margin of the modiolus and stria vascularis) and the tissue lining of the perilymphatic (RWM, margins of ST and SV) and endolymphatic (scala media) spaces (Figure 1, supplementary Videos 1, 2, and 3).

**Figure 1.**
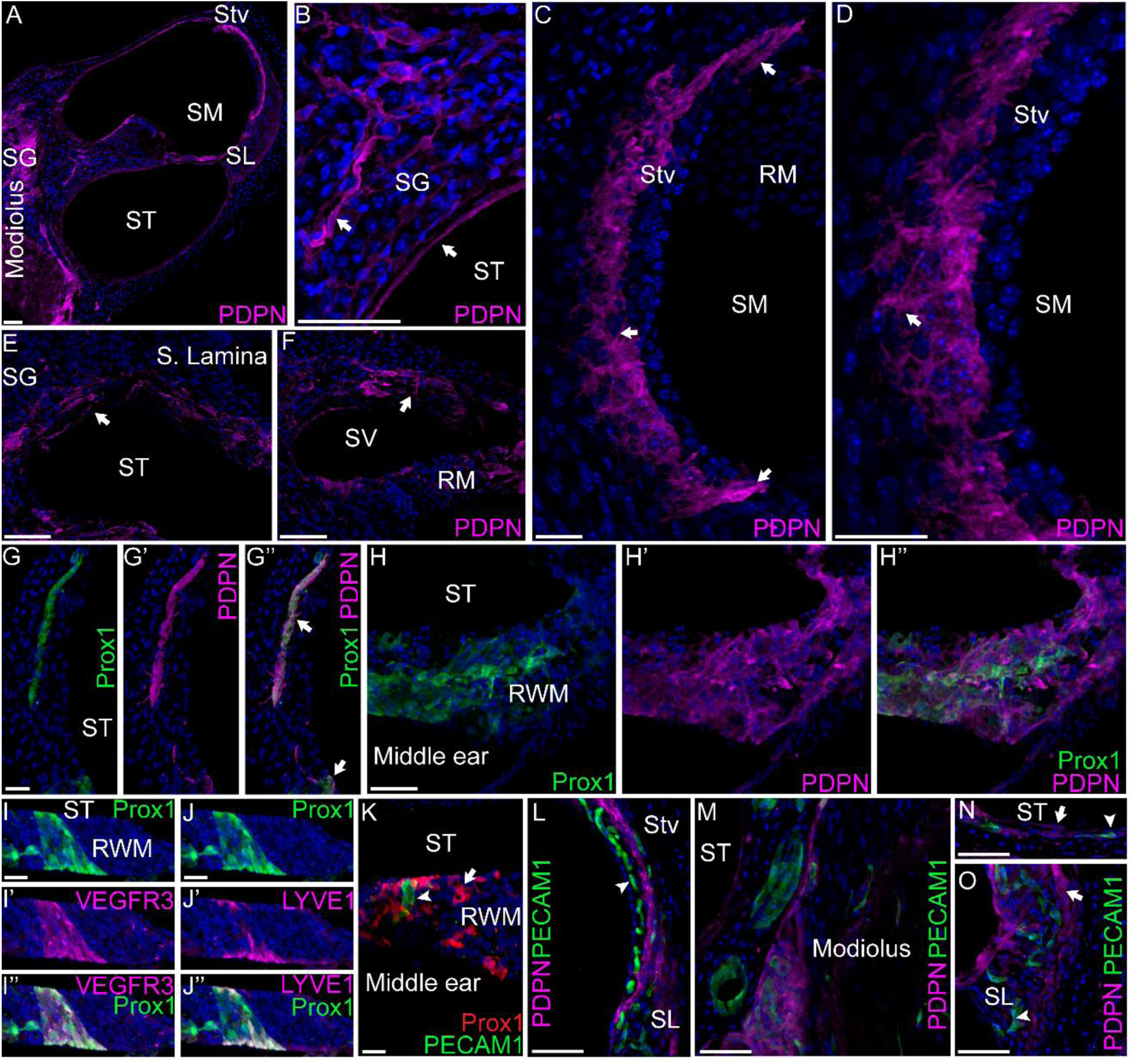
Identification and spatial organization of the lymphatic network in mouse cochlear sections. Identification and determination of the spatial organization of cochlear lymphatics were performed in 8–12-week-old C57BL/6 and Prox1-GFP transgenic reporter mice on a C57BL/6 background. Differentiation of lymphatics from blood vessels was done using a PECAM1-GFP reporter mouse. **A-B**. Representative images of 30μm, and **C-F.** 100 μm thick cochlear sections parallel to the midmodiolar plane from mouse cochlea labeled with anti-podoplanin (PDPN) antibody. **A**. PDPN+ vascular structures in stria vascularis (Stv), spiral ligament (SL), spiral ganglion (SG),(64) and modiolus (M). **B**. PDPN+ cells in SG and the margin of the ST. **C**. PDPN+ cells in the stria vascularis. In addition to the smaller PDPN+ vascular structures throughout the stria vascularis, the upper and lower ends of the stria vascularis harbor larger PDPN+ vessels. D. Magnified view of the stria vascularis showing PDPN+ vessels. E. PDPN+ vascular structures along the margin of ST and F. SV. **G-J.** Sections from Prox-GFP reporter mice labeled with anti-GFP and anti-PDPN antibodies. **G**, **G’,** and **G’’**. 30μm cochlear sections parallel to midmodiolar plane from Prox1-GFP reporter mice were labeled with anti-PDPN antibody. **G**. Prox1 expression, **G’**. PDPN expression, **G’’** Colocalization of Prox1 and PDPN in vascular structure adjacent to the margin of the ST. **H, H’,** and **H’’**.100μm sections of round window membrane (RWM) parallel to midmodiolar plane from Prox1-GFP reporter mice were labeled with anti-PDPN antibody. **H**. Prox1 expression, **H’** PDPN expression, and **H’’** colocalization of Prox1 and PDPN in the RWM. Prox1 and PDPN expression are observed on the perilymphatic (inner ear) and the middle ear side of the RWM. Prox1 and PDPN expression are also observed to be extended to the marginal cells of the ST. **I, I’,** and **I’’**. 60μm sections of RWM parallel to midmodiolar plane from Prox1-GFP reporter mice were labeled with anti-VEGFR3 antibody. **I**. Prox1 expression, **I’** VEGFR3 expression, and **I’’**. colocalization of Prox1 and VEGFR3 in the RWM. **J, J’,** and **J’’**. 60μm sections of round window membrane parallel to midmodiolar plane from Prox1-GFP reporter mice were labeled with anti-LYVE1 antibody. **J**. Prox1 expression, **J’** LYVE1 expression, and **J’’** colocalization of Prox1 and LYVE1 in the round window membrane. **K.** PECAM1-GPF and Prox1-tdTomato mice were crossed, generating blood vessel-lymphatic dual reporter (Prox1-tdTomato: PECAM1-GFP) mice. Representative image of a 100μm thick section of the round window membrane parallel to the midmodiolar plane shows Prox1+ lymphatic vasculature distinct from PECAM1+ blood vessels. **L-O.** 30μm thick sections from blood vessel reporter (PECAM1-GFP) mice were labeled with anti-PDPN antibody. PDPN+ lymphatics are distinct from PECAM1+ blood vessels in **L**. Stria vascularis, **M**. Modiolus, **N.** Margin of the ST, and **O**. Spiral ligament. Spiral ligament shows a PDPN+ vascular network along the outer margin. Arrows: PDPN+ lymphatic vessels, and arrowheads: PECAM1+ blood vessels. SG: spiral ganglion, ST: scala tympani, SM: scala media, SV: scala vestibuli, Stv: stria vascularis, RM: Reissner’s membrane, RWM: round window membrane. SL: spiral lamina. In all cases, Hoechst labeling is shown in blue. Scale bars represent 50μm. n=3 for each condition. Arrows indicate lymphatics, arrowheads indicate blood vessels.

#### 2.1.2. Cochlear lymphatics in whole mounts

Labeling of mid-modiolar sections provided a global overview of the spatial organization of the lymphatic system in the cochlea. Next, we immunolabeled whole-mount cochlear preparations to more readily examine lymphatic distribution in the organ of Corti and the bony spiral lamina. To distinguish the lymphatic vessels from SGNs and their peripheral processes, we used whole mounts from Thy1-YFP mice and labeled them with anti-PDPN antibody. In the bony spiral lamina, PDPN+ lymphatic vessels, distinct from Thy1+ SGN peripheral processes, were identified (Figure 2 and Supplementary Video 4). Within the organ of Corti, PDPN+ lymphatic vessels were seen underneath myosin VIIa+ hair cells (Figure 2 and Supplementary Video 5).

**Figure 2.**
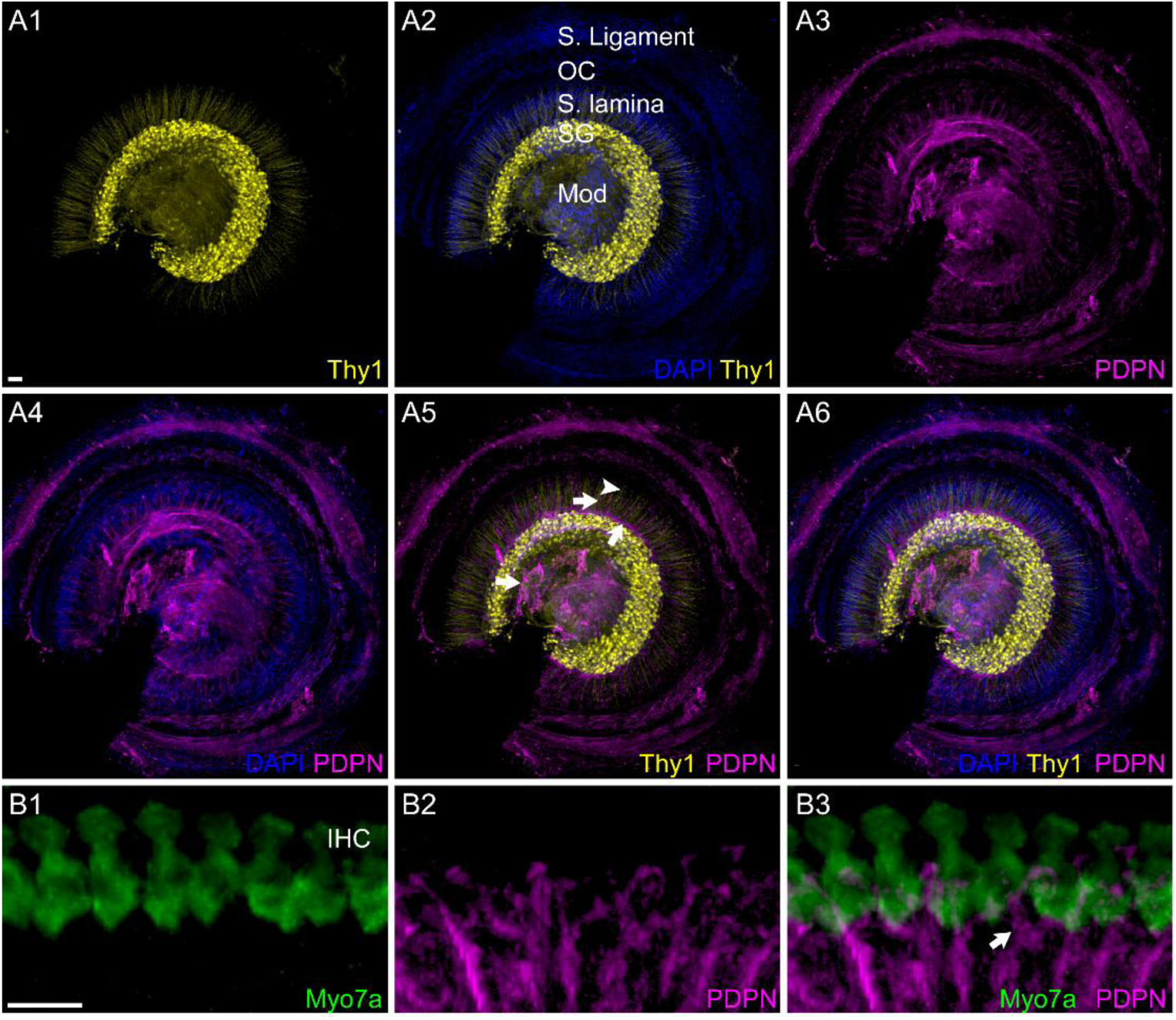
Identification and determination of the spatial organization of the cochlear lymphatic network in mouse cochlear whole mounts. A1-A6. Whole mounts from Thy1^+/YFP^ (neuron reporter) mice were labeled with PDPN and Hoechst 3332. **A1.** Thy1 expression in the whole mount showing Thy1 expression in both SGN soma and peripheral processes. **A2.** Thy1 and Hoechst labeling identify different areas of the cochlea (Mod: modiolus, SG: spiral ganglion, SL: spiral lamina, OC: organ of Corti, S L: spiral ligament). **A3.** PDPN immunolabeling in whole mount. **A4.** Anti-PDPN and Hoechst labeling of whole mount. PDPN+ cells are observed within the modiolus, spiral ganglion, spiral lamina, organ of Corti, and spiral ligament. **A5.** PDPN+ lymphatics are distinct from Thy1+ SGN fibers. PDPN+ vascular structures are shown with arrows, SGN neurites are shown with arrowheads. **A6.** Thy1, PDPN, and Hoechst labeling of a cochlear whole mount preparation. **B1-B3.** Cochlear whole mount labeled with anti-PDPN and anti-Myosin 7a (hair cell). **B1.** Myosin 7a labeling shows a row of inner hair cells. **B2.** PDPN immunolabeling shows PDPN+ lymphatics. **B3.** Merge of Myosin 7a and PDPN labeling, showing PDPN+ lymphatics in the proximity of the base of hair cells. Scale bars: 50 μm. n=3 for each condition.

### 2.2. Spatial organization of MHCII+ antigen-presenting cells and CD11c+ dendritic cells in adult mouse cochlea mirrors the spatial organization of the cochlear lymphatic network

MHC II-mediated antigen presentation is a crucial step in the immune surveillance system, with dendritic cells serving as the primary APCs.(10) In healthy organs, including the brain, APCs are commonly found in close spatial relationship to the lymphatic network.(21) Undamaged CX3CR1^+/GFP^ mouse cochleae labeled with anti-MHCII antibody revealed that CX3CR1+MHCII+ cells are distributed throughout the cochlea, near the vascular planes (outer margin of the modiolus) and along the margin of the perilymphatic spaces (RWM, margins of ST and SV) and spiral ganglion. The distribution of MHCII+ cells in the cochlea mirrors that of the cochlear lymphatic network, except in the stria vascularis, where we identified an extensive lymphatic network but no significant MHCII expression (Figure 3). In the CD11c-DTR/GFP dendritic cell reporter mice, CD11c+ cells are distributed throughout the cochlea, near the vascular planes (outer margin of the modiolus, stria vascularis), and along the margin of the perilymphatic spaces (RWM, margins of ST and SV (Figure 3). While CX3CR1+MHCII+APCs are not found in the stria vascularis, a population of CD11c+ cells is seen in the stria vascularis. Overall, the cumulative distribution of CX3CR1+MHCII+APCs and CD11c+ dendritic cells mirrors that of the cochlear lymphatic vascular network.

**Figure 3.**
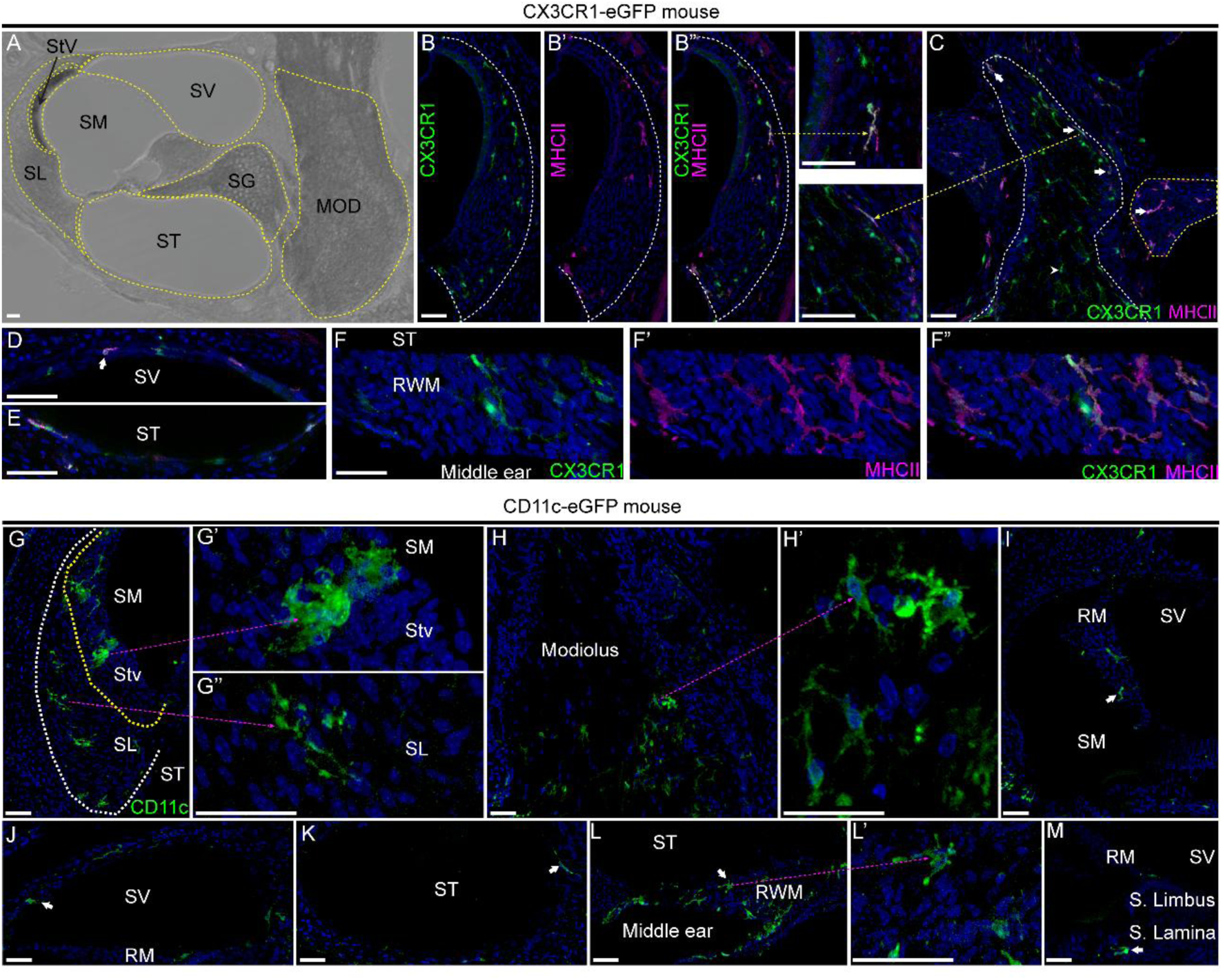
Spatial Organization of the MHCII+ and CD11c cells in the cochlea. Spatial organization of MHCII+ antigen-presenting cells (APCs) and CD11c+ dendritic cells was determined in two transgenic reporter mouse models: CX3CR1^+/GFP^ is a reporter for mononuclear phagocytes (MNPs), while CD11cDTR/GFP is a reporter specific for dendritic cells. All mice used in these experiments are between 8-16 weeks age. **A**. Light microscopic image of a 30 µm-thick section parallel to the midmodiolar plane of the cochlea showing different areas within the cochlea outlined by broken yellow lines (SL: spiral ligament, Stv: stria vascularis, SG: spiral ganglion, MOD: Modiolus, SV: scala vestibuli, SM: scala media, ST: scala tympani). **B-F’’.** 30 µm-thick sections parallel to the midmodiolar plane of the cochlea from CX3CR1^+/GFP^ mice were labeled with anti-MHCII antibody. **B, B’, B’’**. MHC II labeling of the spiral ligament of CX3CR1^+/GFP^ mice. **B**. CX3CR1+ mononuclear phagocytes in the spiral ligament in the lateral wall of the base of the cochlea, **B’** MHCII+ expression in the spiral ligament in the lateral wall of the base of the cochlea. **B’’.** Merge of **B** and **B’**. CX3CR1+MHCII+ mononuclear phagocytes are located near the outer margins of the spiral ligament. The broken yellow arrow shows a higher magnification of the CX3CR1+MHCII+ mononuclear phagocyte. **C**. Section showing modiolus (outlined by broken white line) and spiral ganglion (outlined by broken yellow line). CX3CR1+MHCII+ cells are located near the margin of the modiolus, where the epineurium of the cochlear nerve is located. The broken yellow arrow points towards a higher magnification image of the CX3CR1+MHCII+ mononuclear phagocyte. Within the spiral ganglion, nearly all the CX3CR1+ cells are MHCII+. **D**. ST **E**. Scala vestibuli showing CX3CR1+MHCII+ mononuclear phagocytes along the margin. **F, F’, F’’**. 100 µm-thick sections parallel to the midmodiolar plane that includes the RWM from CX3CR1^+/GFP^ were labeled with anti-MHCII antibody. **F**. CX3CR1+ PMN cells in the RWM. **F’.** MHCII expression in the RWM . **F’’.** Merge of **F and F’** showing CX3CR1+MHCII+mononuclear phagocytes in the RWM (n=3), arrow points towards MHCII+ cells. **G-M.** 100 µm-thick sections parallel to the midmodiolar plane from the CD11c-DTR/GFP dendritic cell reporter mice cochleae were labeled with anti-GFP antibody. **G, G’, G’’.** CD11c+ dendritic cells in the lateral wall of the base of the cochlea, which includes the spiral ligament (SL) and stria vascularis (Stv). **G.** Spatial organization of CD11c+ dendritic cells in the lateral wall of the base of the cochlea: along the outer margin of the spiral ligament and in the stria vascularis. CD11c+ cells in the cochlea have distinct morphology in different locations. **G’.** Magnified image showing a large CD11c+ cell in the stria vascularis with a large cell body and few cell processes. **G’’.** Magnified image showing a CD11c+ cell in the spiral ligament with a small cell body and numerous cell processes. H, H’. CD11c+ dendritic cells in the modiolus of the cochlea. H. CD11c+ dendritic cells along the outer border of the modiolus of the cochlea. H’. Magnified images of the CD11c+ dendritic cells from the modiolus of the cochlea. CD11c+ dendritic cells in the **I.** Reisner’s membrane, **J.** margin of scala vestibuli, **K.** margin of ST, **L, L’.** RWM and **M.** bony spiral lamina. SM: scala media, RM: Ressiner’s membrane, SL: spiral limbus, SL: spiral lamina. Hoechst 3332 labeling shown in blue in all images. Scale bars: 50 μm. n=3 for each condition.

### 2.3. The round window membrane is a niche for an ‘immune-surveillance system’ harboring antigen-presenting cells and lymphatics

As insertion of electrode arrays through the round window membrane (RWM) results in reduced trauma and fibrosis compared to cochleostomy or extended round window insertion, most surgeons favor the round window approach for cochlear implant surgery when feasible.(8) We found that RWM harbors a rich network of blood vessels, lymphatics (Figure 1), MHCII+APCs, and dendritic cells (Figure 3). We further examined the spatial relationships between the lymphatics and MCHII+APCs in the RWM. Within the RWM, the PDPN+ lymphatic network and MHCII+APCs were found in close proximity (Figure 4).

**Figure 4.**
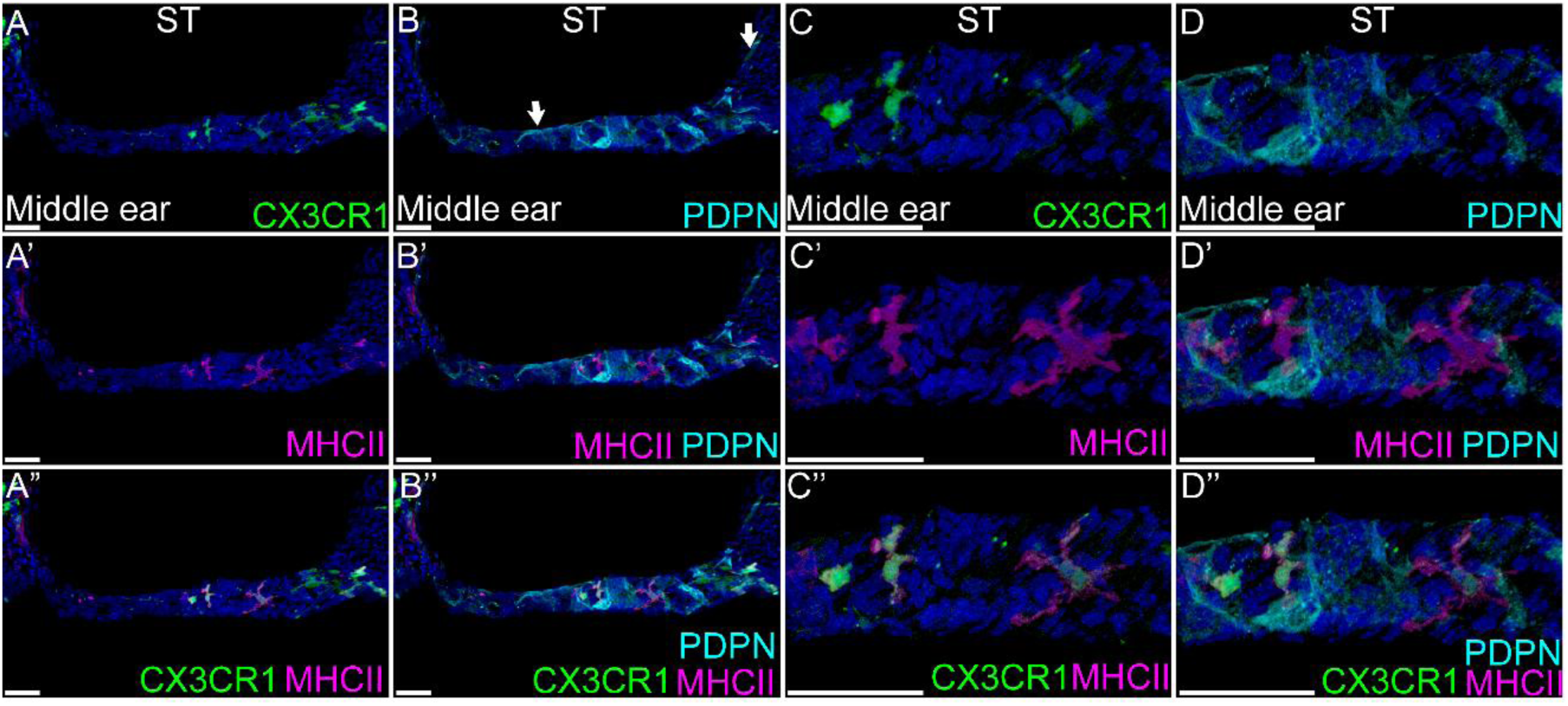
Lymphatics and MHCII+ APCs in the round window membrane. 100 µm-thick sections parallel to the midmodiolar plane that include the RWM from CX3CR1^+/GFP^ were labeled with anti-MHCII and anti-PDPN antibody. **A-A’’ and B-B’’.** Low magnification image of the RWM. **A.** CX3CR1 expression in the RWM, **A’.** MHCII expression in the RWM. **A’’.** Merge of A and A’ showing CX3CR1+MHCII+ cells in RWM. **B.** PDPN expressing lymphatics in RWM. A network of lymphatics is observed in the RWM. Arrows indicate lymphatics in the ST side of the round window and the margin of the ST adjacent to the round window. **B’.** PDPN and MHCII expression in RWM. **B’’.** PDPN, CX3CR1, and MHCII expression in RWM. CX3CR1+MHCII+ APCs are in proximity to PDPN+ lymphatics within the RWM. **C-C’’** and **D-D’’**. Magnified images of the RWM showing the morphology of MHCII+APCs and PDPN+ lymphatics. **C.** CX3CR1 expression in RWM. **C’.** MHCII expression in RWM. **C’’.** Merge of CX3CR1 and **C** and **C’** showing CX3CR1+MHCII+ APCs within the RWM with numerous processes. **D.** PDPN expressing lymphatic vessels forming a lymphatic network in the RWM. **D’.** MHCII+ cells are in proximity to the PDPN+ lymphatic network in the RWM. **D’’.** CX3CR1+MHCII+ APCs and PDPN+ lymphatics are closely associated in RWM.

### 2.4. MHCII+ antigen-presenting cells and CD11c+ dendritic cells are recruited in the implanted mouse cochlea

#### 2.4.1. Increased density of CX3CR1+MHCII+ antigen-presenting cells in implanted mouse cochlea

We previously described that following implantation of electrically active cochlear implants, CX3CR1+ MHCII+ cells are recruited in the implanted cochlea.(22) Following implantation of inactive cochlear implants in CX3CR1^+/GFP^ mice, an increased density of CX3CR1+MHCII+ antigen-presenting mononuclear phagocytes was likewise seen in the ST of the implanted cochlea compared to the contralateral unimplanted cochlea (p=0.0827, paired t-test, n=4) (Figure 5). Supplementary information and Supplementary Figure 5 further describe recruitment of MHCII+ APCs in other regions of the cochlea following implantation.

**Figure 5.**
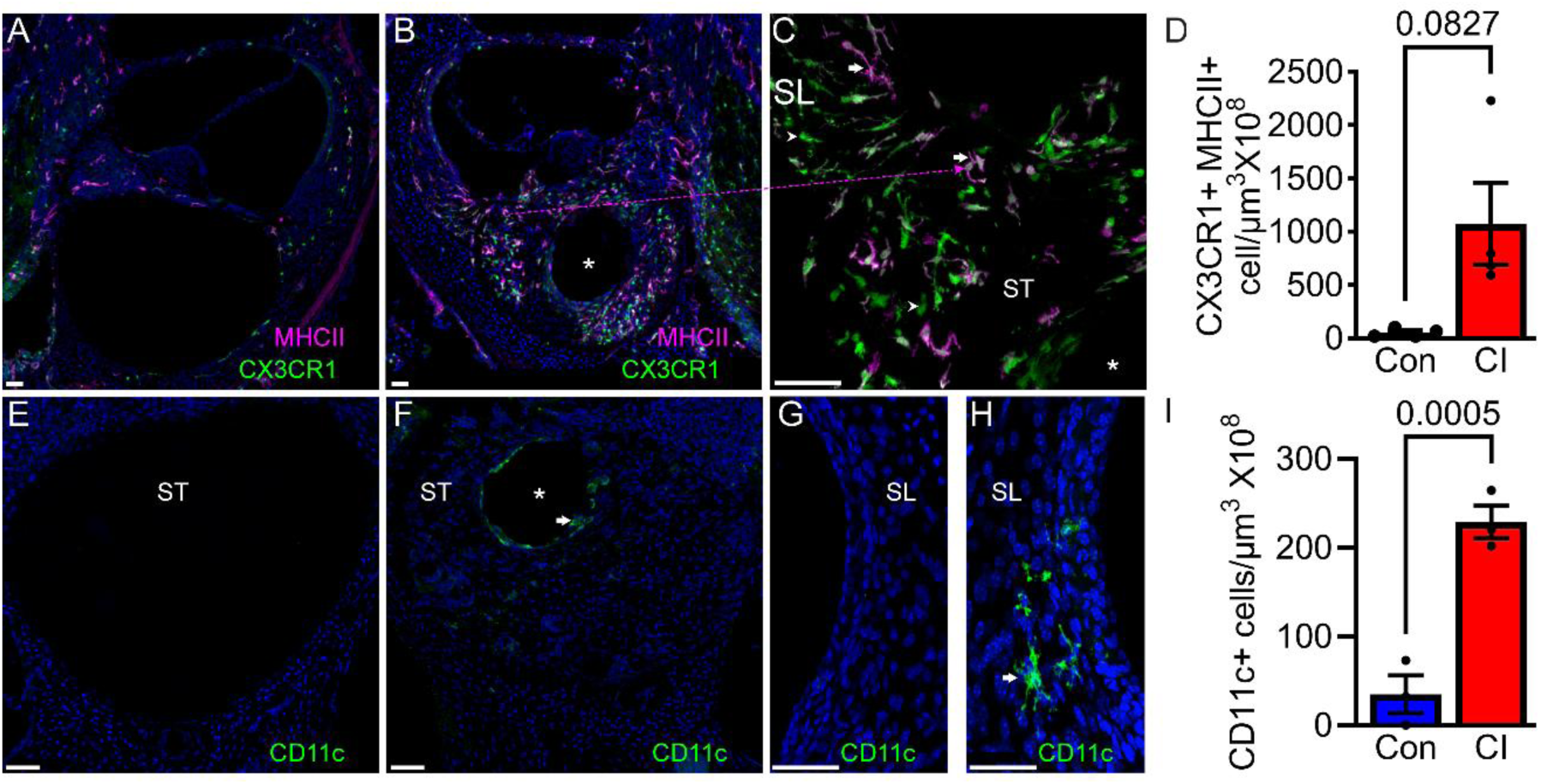
MHCII+APC and CD11c+ dendritic cells respond to cochlear implantation. **A-D.** 28 days after cochlear implantation in CX3CR1^+/GFP^ reporter, 30 µm-thick sections parallel to the midmodiolar plane of the cochlea were labeled with anti-MHCII antibody. **A**. Unimplanted (contralateral) cochlea. **B**. Implanted cochlea showing increased density of CX3CR1+MHCII+ mononuclear phagocytes in ST of the base of the cochlea and the spiral ligament. **C.** Magnified images from an implanted cochlea showing both CX3CR1+ MHCII+ APCs (arrows) and CX3CR1+MHCII-(arrowheads) MNPs. **D.** Quantitative analysis of CX3CR1+MHCII+ mononuclear phagocytes is shown for the ST. Statistical analyses were performed using paired t-test. n=4 for each group. Error bar indicates SEM. Statistical analyses with paired t-test. p-value was shown on each graph; a p-value of <0.05 was considered statistically significant. **E-I.** 28 days after cochlear implant surgeries in CD11c-DTR/GFP reporter, 30 µm-thick sections parallel to the midmodiolar plane of the cochlea were labeled with anti-GFP antibody. Representative images of the ST of the cochlea from a CD11c-DTR/GFP mouse. **E**. Contralateral (unimplanted) cochlea, and **F**. Implanted cochlea. Representative images of the spiral ligament of the cochlea from CD11c-DTR/GFP mouse. **G.** Contralateral (unimplanted) and **H**. Implanted cochlea. Arrow indicates CD11c+ dendritic cells adjacent to the implant tract (asterisk) **I.** Quantitative analysis of CD11c+ dendritic cells for the ST. Scale bars: 50 µm. Statistical analyses were performed using paired t-test. n=3 for each group. Error bar indicates SEM. p-value was shown on each graph; a p-value of <0.05 was considered statistically significant.

#### 2.4.2. Increased density of CD11c+ dendritic cells in the implanted mouse cochlea

In a CD11c-DTR/GFP mouse model, an increased density of CD11c+ dendritic cells was evident in the ST of the implanted cochlea compared to the unimplanted contralateral cochlea (p=0.0005, paired t-test, n=3) (Figure 5). Notably, CD11c+ dendritic cells reside adjacent to the implant tracts in the ST (asterisk, Figure 5), suggesting that they respond to the surface properties of the implanted biomaterials. Recruitment of CD11c+ dendritic cells following implantation in other cochlear areas is described in the supplementary information and Supplementary Figure 5.

### 2.5. Lymphangiogenesis within the scala tympani following cochlear implantation, trafficking dendritic cells

#### 2.5.1. Lymphangiogenesis within the scala tympani following cochlear implantation

The recruitment of MHCII+APCs and CD11c+ dendritic cells within the cavity of the ST raised the question of whether these APCs are trafficked through lymphatic channels. This is of particular relevance for the dendritic cells, as we noted that CD11c+ dendritic cells localize very close to the implant tract where lymphatic vessels are not present in the lumen of the unimplanted ST. As previously described, lymphatics are distributed in the RWM and along the margin of the ST (Figure 1) in the unimplanted cochlea, which are distant from the implant tract. To address this question of dendritic cell trafficking through the cochlear lymphatics, we implanted lymphatic-dendritic cell dual reporter (Prox1-tdTomato: CD11c-DTR/GFP) mice. Twenty-eight days after cochlear implantation, there was a dramatic increase in density of Prox1+ vascular structures within the ST of the mouse cochlea compared to the contralateral unimplanted cochlea (p=0.0218, paired t-test, n=3) (Figure 6). These Prox1+ cells also expressed other markers of lymphatic endothelial cells (VEGFR3, PDPN, LYVE1), validating their identity as lymphatic vessels (Figure 6). Notably, these newly developed lymphatic vessels within the ST are embedded in the FBR surrounding the implant tract.

**Figure 6.**
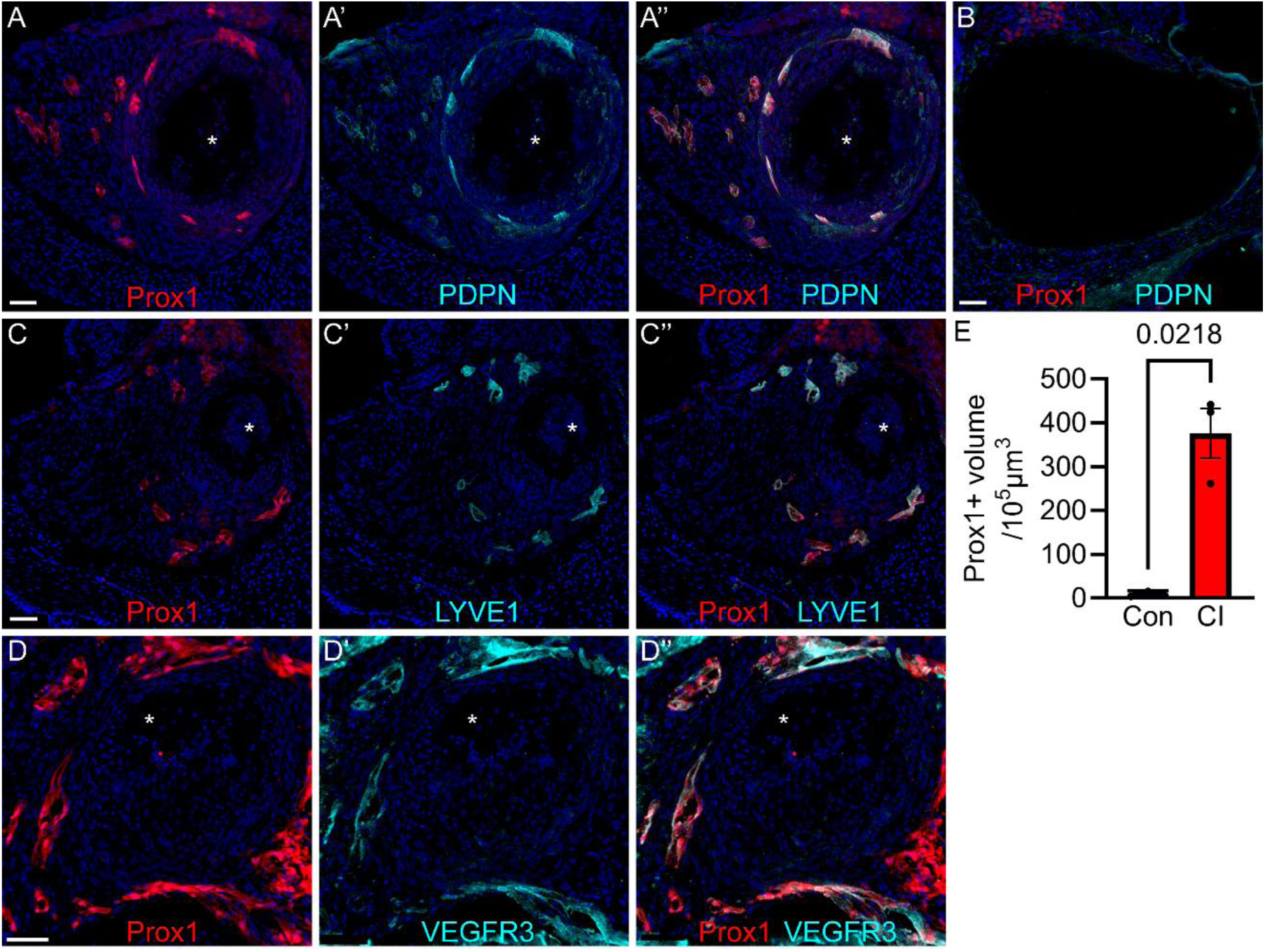
Proliferation of lymphatics within the scala tympani of the cochlea following cochlear implantation. Prox1-tdTomato mice were implanted, and 28 days post-CI, 30μm sections parallel to the midmodiolar plane were labeled with anti-PDPN. **A, A’,** and **A’’**. ST of implanted cochlea. **A**. Prox1-tdTomato, **A’** PDPN, and **A’’** merge of Prox1 and PDPN in the implanted cochlea. Colocalization of Prox1 and PDPN in the implanted cochlea indicates lymphangiogenesis within the ST of the implanted cochlea. **B**. ST of unimplanted, contralateral cochlea. The implanted cochleae were further labeled with LYVE1 **(C, C’, and C’’)**, and VEGFR3 **(D, D’, and D”)** antibodies. **C**. Prox1, **C’** LYVE1, and **C”** merge of Prox1 and LYVE1 showing colocalization of Prox1 and LYVE1. **D**. Prox1, **D’** VEGFR3, and **D”** merge of Prox1 and VEGFR3 showing colocalization of Prox1 and VEGFR3. **E**. Quantification of lymphangiogenesis within the ST of the cochlea. ST periphery was traced, and the volume was measured. Creating a mask for individual channels, the volume of Prox1+ lymphatic labeling was measured, which was then divided by the volume of ST, expressed/10^5^ μm^3^. n=3 for each. Error bar indicates SEM. Statistical analyses with paired t-test. p-value shown on the graph.

#### 2.5.2. Newly developed lymphatics in the implanted mouse cochlea act as conduits for CD11c+ dendritic cells

In implanted Prox1-tdTomato: CD11c-DTR/GFP (lymphatic-dendritic cell dual reporter) cochleae, CD11c+ dendritic cells were detected within the newly formed lymphatic vessels (Figure 7 and Supplementary Video 6). We identified these cells as intralymphatic dendritic cells as previously described.(23) In the ST of the implanted cochlea an increased density of dendritic cells and intralymphatic dendritic cells was noted (p=0.0102, paired t-test, n=3).

**Figure 7:**
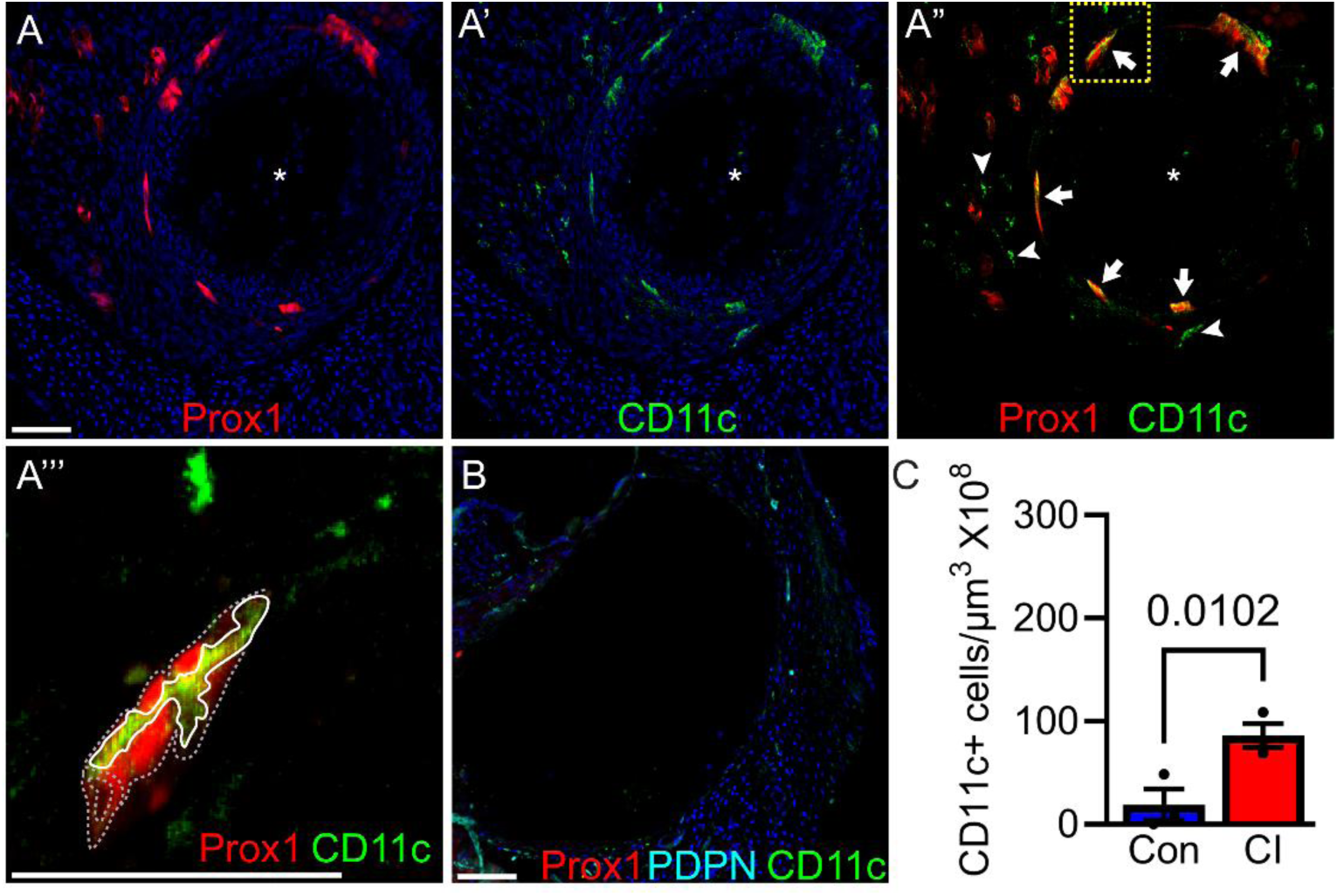
Proliferated lymphatics following cochlear implantation act as conduits for dendritic cells. Prox1-tdTomato: CD11c-DTR/GFP mice were implanted, and 28 days post-CI, 30μm sections parallel to the midmodiolar plane were labeled with anti-GFP antibody. **A, A’, A’’.** Representative images from the implanted Prox1-tdTomato:CD11c-DTR/GFP mouse cochlea. **A**. Prox1+ lymphatic vessel proliferation around the implant tract (asterisk) **A’**. CD11c+ dendritic cells are observed within the ST of the implanted cochlea, and **A”**. CD11c+ dendritic cells are observed within the ST of the implanted cochlea inside the lumen of Prox1+ proliferated lymphatics. Arrows point toward proliferated lymphatics harboring intralymphatic CD11c+ dendritic cells. Arrowheads show extralymphatic dendritic cells. **A’’’.** A magnified version of the inset from A’’. Prox1+ lymphatic vessel has been outlined with a dashed grey line. An intralymphatic CD11+ dendritic cell is outlined using a white continuous line. **B**. Unimplanted contralateral cochlea from Prox1-tdTomato: CD11c-DTR/GFP mice labeled with PDPN. **C**. Quantification of lymphatic trafficking of CD11c+ dendritic cells post-CI. Creating masks for PDPN and CD11c, intralymphatic dendritic cells were identified and quantified/10^8^ μm^3^ of ST. n=3 for each. Error bar indicates SEM. Statistical analyses with paired t-test. p-value was shown on the graph. Con: Contralateral cochlea, CI: Implanted cochlea. Scale bar 50μm.

**Figure 8.**
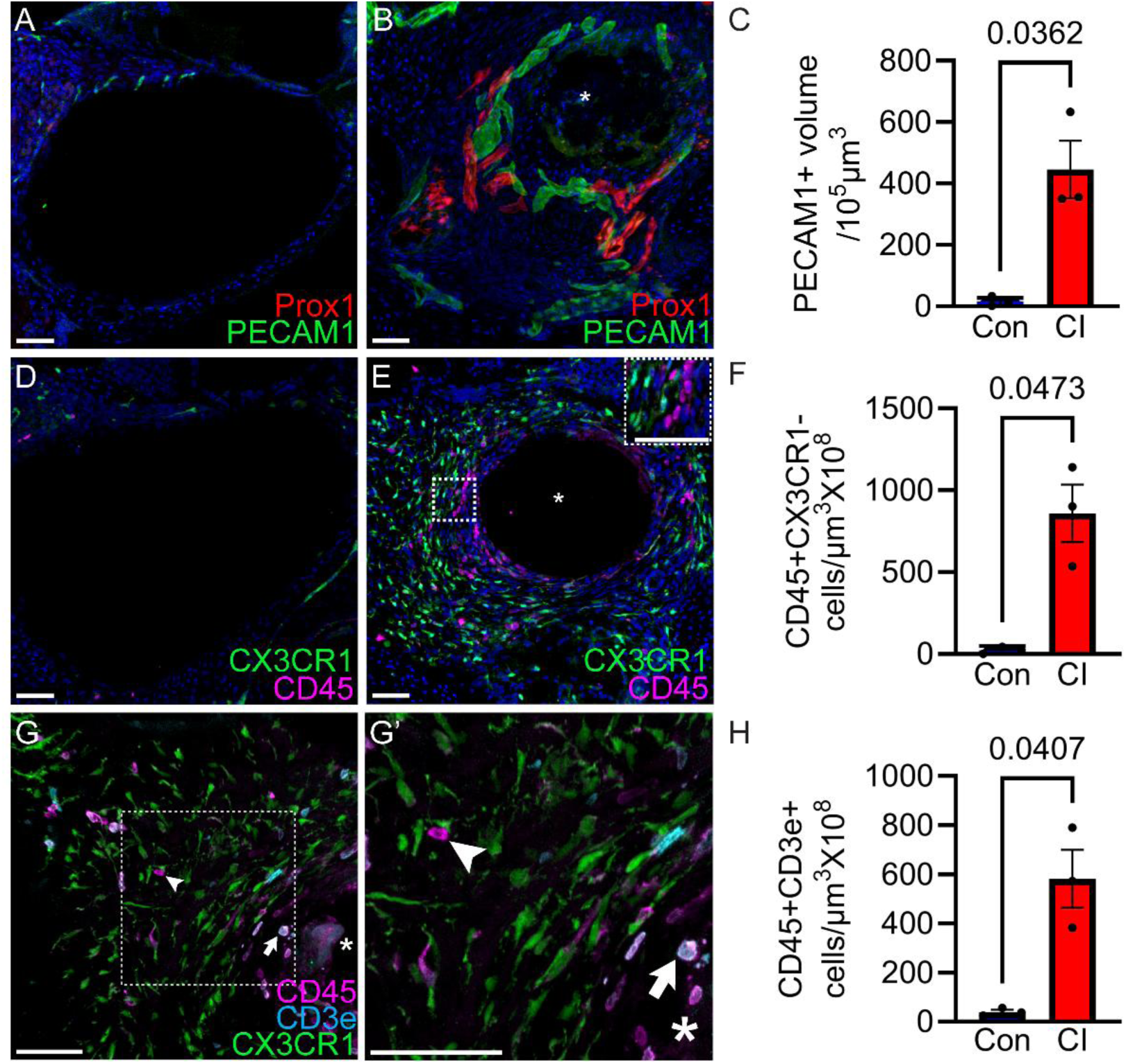
Proliferation of blood vessels and infiltration of blood-derived immune cells within the implanted cochlea. Prox1-tdtomato: PECAM1-GFP (Lymphatic-blood vessel dual reporter) mice were implanted in the left cochlea. 28 days post-CI, 30μm midmodiolar sections were labeled with anti-GFP antibody. **A**. Unimplanted, contralateral (con) shows some blood vessels and lymphatics lining the ST, but not within the ST lumen. **B**. Implanted cochlea showing PECAM1+ blood vessel proliferation around the implant tract (asterisk), distinguished from Prox1+ proliferated lymphatic vessels. **C**. Quantification of blood vessel proliferation in the ST, which was traced manually, and the volume was measured. The volume of PECAM1+ mask was also measured and expressed as /10^5^ μm^3^ of ST. n=3 for each condition. Error bar indicates SEM. Statistical analyses with paired t-test. p-value was shown on the graph. Con: Contralateral cochlea, CI: Implanted cochlea. CI causes dendritic cell infiltration and increases lymphatic trafficking. To identify different immune cell types, including lymphocytes, we implanted CX3CR1^+/GFP^ reporter mice. 28 days post-CI, 30μm sections parallel to the midmodiolar plane were labeled with an antibody against pan-leukocyte marker CD45 to differentiate CX3CR1+ mononuclear phagocytes and other non-mononuclear phagocytic (CX3CR1-CD45+) immune cells. **D**. Midmodiolar section of unimplanted, contralateral cochlea. **E**. Implanted side showing mononuclear phagocytes (CX3CR1+) and non-mononuclear phagocytic (CX3CR1-CD45+) cells. The inset shows a magnified view of mononuclear phagocytes (CX3CR1+) and other (CX3CR1-CD45+) immune cells. **F**. Quantification of (CX3CR1-CD45+) immune cells was done by counting the number of cells and dividing by the volume of ST that was traced manually, expressed/10^8^μm^3^ of ST. n=3 for each condition. Error bar indicates SEM. Statistical analyses with Paired t-test. p-value was shown on the graph; a p-value of <0.05 was considered statistically significant. Con: Contralateral cochlea, CI: Implanted cochlea. **G**. Adaptive T lymphocytes were labeled with anti-CD3e antibody. A significant subset of CX3CR1-CD45+ cells in the implanted ST are CD3e+ T cells (arrow) with some CX3CR1-CD45+CD3e- cells (arrowhead). **G’.** Shows a magnified view of the inset from G. **H.** Quantification of (CX3CR1-CD45+CD3e+) immune cells. n=3 for each condition. Error bar indicates SEM. Statistical analyses with paired t-test. p-value was shown on the graph. Con: Contralateral cochlea, CI: Implanted cochlea. Scale bar 50μm.

### 2.6. Angiogenesis and infiltration of blood-derived immune cells in the implanted mouse cochlea

#### 2.6.1. Angiogenesis in the implanted mouse cochlea

Dendritic cells represent blood-derived immune cells.(24) While blood vessels exist around the perilymphatic space, they are limited to the RWM and along the margins of the ST and SV (Figure 1). We identified dendritic cells within the cavity of the implanted cochlea close to the implant tract (Figure 5), distant from the nearest blood vessels, and in the RWM and the margin of the ST in an unimplanted cochlea. Moreover, other blood-derived immune cells (e.g., T cells, B cells) have been reported in the implanted cochlea.(8) How blood-derived immune cells access the inflammatory response surrounding the implant tract is unknown.

Angiogenesis has been reported in the FBR to implanted biomaterials in other tissues, potentially leading to immune infiltration.(25) We implanted Prox1-tdTomato: PECAM1-GFP (lymphatic-blood vessel dual reporter mice) to differentiate angiogenesis from lymphangiogenesis. As shown in Figure 7, 28 days following cochlear implantation, a striking new growth of PECAM1+ blood vessels emerges within the ST of the base of the cochlea around the implant tract (p=0.0362, paired t-test, n=3), supporting that angiogenesis may facilitate immune cell infiltration during the FBR.

#### 2.6.2. Blood-derived immune cells, including T lymphocytes, infiltrate into the implanted cochlea

As angiogenesis is robust in the implanted ST, we asked the extent to which blood-derived immune cells infiltrate the implanted cochlea. Mononuclear phagocytes (MNPs) are classified as either tissue-resident or infiltrating(26); dendritic cells are a type of infiltrating (blood-derived) MNP that we observed in the implanted cochlea. Non-MNP immune cells, on the other hand, are blood-borne, and derived from bone marrow(27). To identify non-MNP immune cells, we implanted CX3CR1^+/GFP^ reporter mice and labeled the harvested cochlea with anti-CD45 (pan leukocyte marker) antibody 28 days after cochlear implantation. In the implanted cochlea, CX3CR1-CD45+ non-macrophage immune cell density is higher in ST of the base of the cochlea, compared to the contralateral unimplanted control cochlea (8, 22, 28) (Figure 7). CD3e+ T lymphocyte density is also increased in the ST of the base of the cochlea (p=0.0407, paired t-test, n=3) (Figure 3). Infiltration of non-MNP immune cells in other areas of the cochlea is shown in the supplementary information and Supplementary Figure 6.

### 2.7. Angiogenesis and lymphangiogenesis within the scala tympani of the human cochlea

To detect possible changes in the blood vessels and lymphatic network in the implanted cochlea in humans, temporal bone sections from an 81-year-old male who underwent right cochlear implantation via cochleostomy with full insertion of a straight electrode one year prior to death were collected. Both the implanted and contralateral, unimplanted ears were immunolabeled with markers for blood vessels (CD34) and lymphatics (Lyve1). Proliferation of both blood vessels and lymphatics was evident within the scala tympani of the implanted cochlea (Figure 9).

**Figure 9.**
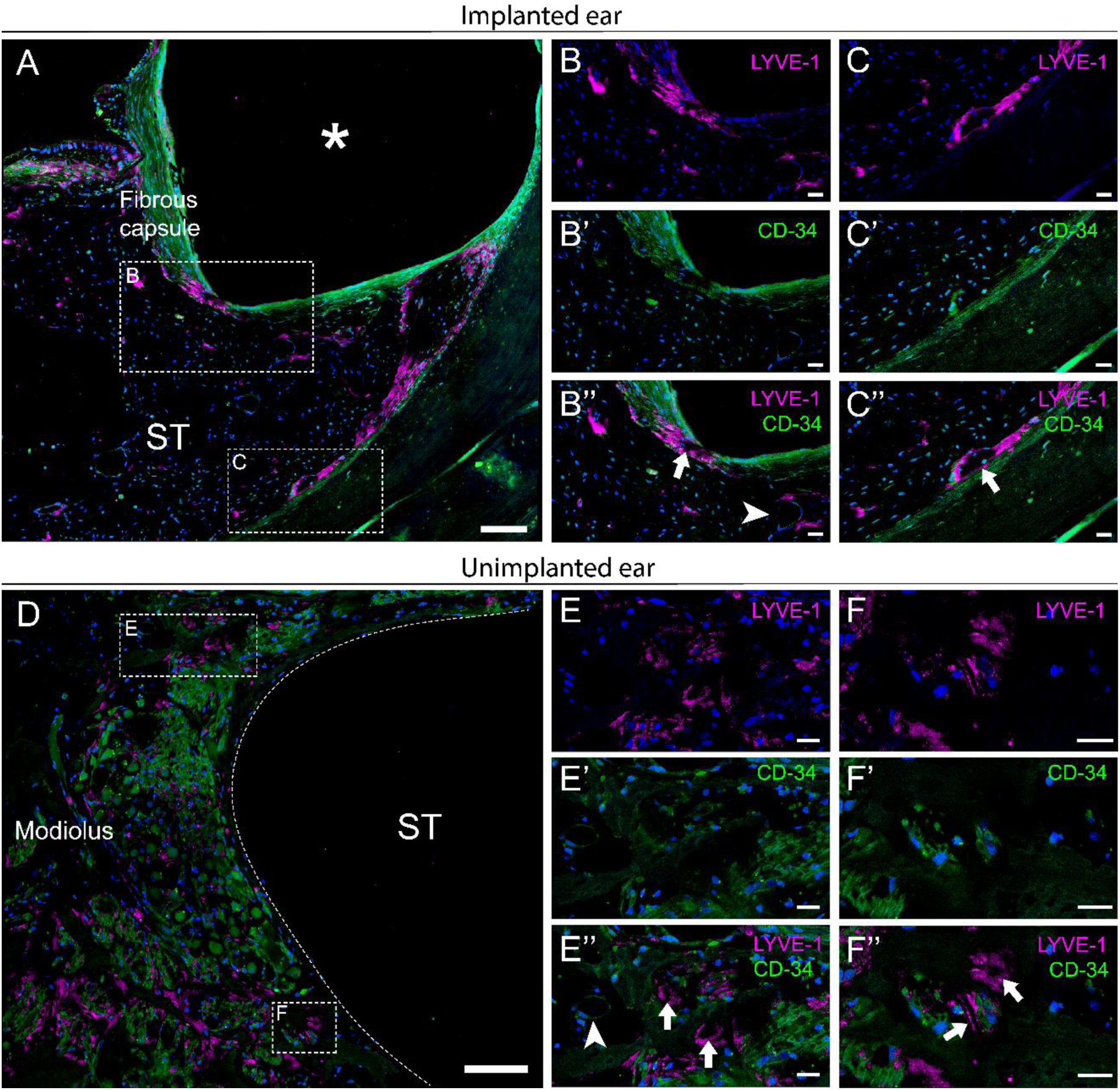
Lymphatic vessels in human cochleae. Immunofluorescence of archival temporal bone sections from an 81-year-old man with progressive bilateral adult-onset sensorineural hearing loss, unilaterally implanted 1 year prior to death with a straight electrode CI. Anti-LYVE-1 for lymphatic vessels (arrowheads), anti-CD34 distinguishes blood vessels (arrows), and Hoechst for nuclear staining. **A,B,C.** Implanted right cochlea shows lymphatic channels are in the fibrosis surrounding the electrode array (asterisk) in the scala tympani (ST) (**A**), including within the fibrotic capsule (**B, B’, B’’**) of the electrode array and at the lateral cochlear wall within the ST (**C, C’, C’’**). **D,E,F.** Contralateral left unimplanted cochlea from the same donor has no fibrosis in the ST and shows lymphatics around the SGN peripheral axons (**E, E’, E’’**), and at the medial wall of the scala tympani around SGN central axons (**F, F’, F’’**), with distinct vessel lumens, similar to mouse cochlea. Scale bars in A, D = 100 um, B-C, E-F = 20 um.

## 3. Discussion

In this study, we provide evidence for a rich network of lymphatic vessels expressing multiple molecular markers within the undamaged mouse cochlea. This lymphatic network extends through most regions of the cochlea, particularly near the RWM, the lining of perilymphatic and endolymphatic spaces, the lateral wall, the spiral ganglion, and along the basilar membrane just below the sensory hair cells.

For many years, the central nervous system (CNS) was considered immune-privileged, partly attributed to a lack of draining lymphatics.(29) Despite evidence from decades ago suggesting that meningeal-based lymphatics could participate in the drainage of cerebrospinal fluid (CSF),(30) molecular marker-based compelling evidence for meningeal lymphatics only emerged relatively recently.(21, 31) Shortly afterward, lymphatics were described as the major draining pathway for CSF.(32) Importantly, the subarachnoid space containing CSF connects to the perilymphatic space in the cochlea via the cochlear aqueduct.(33) Therefore, a major question we addressed was whether cochlear lymphatic vessels are positioned to participate in the drainage of the perilymphatic space Previously, injection of a tracer, keyhole limpet hemocyanin, into the basal turn of the guinea pig cochlea resulted in the labeling of marginal cells of ST, cervical lymph nodes, and spleen, strongly suggesting a connection between the perilymphatic space and the lymphatic system.(18) Moreover, cochlear implant biomaterial (platinum) has been detected in the kidneys of implanted animals(34), implying a draining pathway for perilymphatic fluid and cochlear immune cells. Despite indirect evidence, molecular marker-based evidence for the presence of cochlear lymphatic vessels draining perilymphatic space has been lacking.

Lymphatic vessels have been described in the peripheral(35) and cranial nerves (e.g., optic,(36) trigeminal, glossopharyngeal, vagus, and accessory)(26)(37) in human and animal models.(31) Here, we present evidence for a robust cochlear lymphatic network. Previously, lymphatic vessels have been described in the round window of the cochleae of diseased human ears.(20, 38) Likewise, we observed lymphatics in the round window and also along the margin of the ST in mice without any known pathology. The distribution of lymphatic vessels in the margins of the cochlear perilymphatic spaces (RWM, margins of ST and SV) places them in a strategic location for drainage of perilymphatic spaces. Further, we observed MCHII+ APCs, and CD11c+ dendritic cells and blood vessels near the lymphatic vessels, suggesting the presence of an ‘immune-surveillance system’ adjacent to the perilymphatic spaces.

Cochlear implantation within the perilymphatic space (ST) resulted in the recruitment of the cellular components of the cochlear ‘immune surveillance system’: immune cells, lymphatics, blood vessels, MCHII+ APCs, and CD11c+ cells were recruited around the implant tract. Moreover, lymphatic vessels proliferated around the implant tract, trafficking the dendritic cells. Concurrently, blood vessels proliferated in the ST of the implanted cochlea, potentially playing a key role in the infiltration of blood-derived immune cells. The composite of our data implies that following cochlear implantation, cellular components of the ‘immune surveillance system’ initiate a two-way trafficking of immune cells in the cochlea through proliferated blood and lymphatic vessels.

As the discovery of lymphatic vessels in the nervous system is relatively recent, knowledge of lymphatic proliferation in response to implanted biomaterials is relatively limited. Meningeal lymphatic proliferation with increased lymphatic drainage was observed near chronically implanted electroencephalogram electrodes.(39) Likewise, we find that cochlear implantation results in lymphatic proliferation adjacent to the implanted biomaterials.

Little is known about the role of dendritic cells in the FBR to implanted biomaterials, and to our knowledge, recruitment of dendritic cells in the implanted cochlea has not been reported. Upon recognition of pathogen-associated molecular patterns or damage-associated molecular patterns, dendritic cells migrate to draining lymph nodes, activate T lymphocytes, and initiate adaptive immune responses characterized by immunological memory.(40) Recent studies suggest that dendritic cells play an important role in the host response to biomaterials, including signaling pathways leading to immune activation.(40) The specific role of dendritic cells in the implanted cochlea and dendritic cell trafficking through the lymphatics requires further studies. Interestingly, recent data from patients undergoing sequential cochlear implantation revealed that the second implanted ear had higher electrode impedances and a higher risk of loss of residual hearing, suggesting the possibility of priming of a systemic immune response from prior cochlear implantation.(41, 42) A similar priming of the systemic immune response is seen in sympathetic ophthalmia, where penetrating trauma or surgery to one eye initiates a bilateral, granulomatous inflammation.(43)

Angiogenesis is a crucial component of the FBR.(44) , its significance is unclear. The implanted mouse cochlea is infiltrated by blood-derived immune cells, including T lymphocytes and dendritic cells, similar to the FBR in many organs.(45) We questioned how immune cells enter the cavity of ST, which harbors blood vessels only along the margin.(16) One potential source of post-CI lymphangiogenesis and angiogenesis in the ST is the reactive proliferation of RWM blood and lymphatic vessels in response to cochlear implant trauma through the RWM, which contains an impressive network of lymphatic and blood vessels. The lymphatic and blood vascular network along the margin of ST is another possible source of angiogenesis and lymphangiogenesis. Similarly, MHCII+ APCs and CD11c+ dendritic cells in the RWM and the margin of the ST likely contribute to the immune response to the indwelling cochlear implant. Our data, together with recent publications, imply the development of a functional, ectopic lymphoid structure in the ST adjacent to the cochlear implant with the formation of new vascular and lymphatic structures and the recruitment of a complex milieu of activated immune cells, including MNPs and T lymphocytes.(8, 22, 46)

The FBR depends on the type and long-term presence of the implanted biomaterials since surgical implantation and subsequent withdrawal of the electrode array fail to generate these responses in general.(47, 48) Moreover, dexamethasone-eluting electrode arrays dramatically reduce this FBR in animal models and electrode impedances (a surrogate for inflammatory/fibrotic tissue) in humans.(8, 22, 28) These notable findings confirm that the immune responses in the cochlea following CI can be modulated, carrying important implications for cochlear injury and repair and the ongoing development of advanced CI technologies.(49–53)

Apart from the immune surveillance system closely associated with the perilymph cavity, which is directly affected by cochlear implantation, we also identified the cellular components of the immune surveillance system in other locations in the cochlea. Particularly interesting in this context is the lymphatic vascular network beneath the hair cells, where these cells form synapses with SGNs. The function of these lymphatics in the organ of Corti, near the synapses, and their contribution to the immune and synaptic functions need to be explored. Likewise, the response of cochlear lymphatics and the immune surveillance system to other inflammatory inner ear conditions like aging, ototoxicity, and noise-damage deserves detailed investigation.(50)

## 4. Materials and Methods

### 4.1 Sex as a biological variable

Both male and female mice were used in this study, reflecting the biological relevance of cochlear lymphatics in the inner ear and the clinical significance of cochlear implantation in both sexes. In the human study, a male cochlear implant recipient’s sample was used. This choice was guided by the availability of samples and based on the presence of lymphatics and changes following cochlear implantation in mice from either sex.

### 4.2 Animals

All the experimental protocols on mice in this study were approved by the University of Iowa Institutional Animal Care and Use Committee. For this study, we used both male and female 8–12-week-old mice on a C57BL/6J background. The following mouse strains were used:

#### 4.2.1. Mononuclear phagocyte reporter (CX3CR1-GFP knock-in/knock-out, CX3CR1^+/GFP^) mice

(B6.129P2(Cg)-*Cx3cr1^tm1Litt^*/J, Strain #:005582, RRID:IMSR_JAX:005582, Jackson Laboratory) express GFP in mononuclear phagocytes under control of the endogenous *Cx3cr1* locus.(54)

#### 4.2.2. Neuron Reporter (Thy1^+/YFP^) mice

B6.Cg-Tg (Thy1-YFP)16Jrs/J, Strain #:003709, RRID: IMSR_JAX:003709, Jackson Laboratory) in which SGNs express YFP.(55)

#### 4.2.3 Lymphatic reporter mice

Prox1 is a transcription factor necessary for the development of the lymphatic system.(56) To define the cochlear lymphatic vessels, two lymphatic reporter mice were used in this study:

##### Prox1-GFP mice

STOCK Tg(Prox1-EGFP) KY221Gsat/Mmucd, RRID: MMRRC_031006-UCD. These mice express EGFP in lymphatic endothelial cells under the control of the endogenous *Prox1* locus.

##### Prox1-tdTomato mice

STOCK Tg(Prox1-tdTomato)TA76Gsat/Mmucd, RRID:MMRRC_036531-UCD. These mice express tdTomato in lymphatic endothelial cells under the control of the endogenous *Prox1* locus. Both Prox1-GFP and Prox1-tdTomato mice were obtained from the Mutant Mouse Resource and Research Center (MMRRC) at the University of California at Davis, an NIH-funded strain repository, and were donated to the MMRRC by Nathaniel Heintz, Ph.D., The Rockefeller University, GENSAT(57)

#### 4.2.4. Dendritic cell (CD11c-DTR/GFP) reporter mice

CD11c is a sensitive and specific marker for dendritic cells (DC).(58) CD11c-DTR/GFP transgenic mice have the CD11c promoter (Itgax) directing expression of a diphtheria toxin receptor-enhanced green fluorescent protein (DTR/EGFP) fusion protein in dendritic cells (DC).

#### 4.2.5. Lymphatics-dendritic cell dual reporter (Prox1-tdTomato:CD11c-DTR/GFP) mice

These mice were generated by crossing Prox1-tdTomato and CD11c-DTR/GFP mice.

#### 4.2.6. Blood vessel endothelial cell reporter (PECAM1-GFP) mice

PECAM1-GFP transgenic mice have the PECAM1 promoter (CD31) directing the expression of enhanced green fluorescent protein.(59) In our experiments, we used these mice as blood vessel endothelial cell reporter mice.

#### 4.2.7. Lymphatic-blood vessel dual reporter (Prox1-tdTomato :PECAM1-GFP mice)

These mice were generated by crossing Prox1-tdTomato and PECAM1-GFP mice.

### 4.3. Genotyping

Genotyping was performed using automated standard PCR of genomic DNA from tail samples, performed by Transnetyx genotyping services (https://www.transnetyx.com/)

### 4.4. Cochlear whole mount preparation

Cochlear wholemount dissection and immunolabeling of the adult mouse cochlea were performed as previously described.(60) A detailed description of the cochlear whole mount preparation is provided in the supplementary materials and methods.

### 4.5. Cochlear implantation in a mouse model

Cochlear implantation was performed in the left ear of mice through a round window approach as previously described.(47) A detailed description of the cochlear implantation method is provided in the supplementary materials and methods.

### 4.6. Immunohistochemistry

Under ketamine (80mg/kg) and xylazine (10mg/kg) anesthesia, transcardial perfusion with phosphate-buffered saline (PBS) followed by 4% paraformaldehyde (PFA) was performed. Cochleae were harvested and fixed with 4% PFA at 4°C in the dark. Cochleae were decalcified in 0.1M EDTA (pH 7.5) for 3 days, cryopreserved using increasing (10-30%) concentrations of sucrose solutions as we have described previously, and embedded in O.C.T. as we previously described.(61) 30µm or 60µm thick sections parallel to the mid-modiolar plane were washed with ’washing buffer’ containing Tris-buffered saline (TBS), 0.03% Triton X-100, and 0.1% Tween-20, followed by permeabilization and blocking of samples in ’blocking buffer’ (0.03% Triton X-100 and 0.1% Tween-20, 1% bovine serum albumin in TBS for 2 hours. After blocking and permeabilization, sections were incubated in primary antibodies in ’blocking buffer’ overnight (∼16 h) at 4°C. Following primary antibody application, sections were washed (3 x 5 min) in ’washing buffer’, followed by incubation in blocking buffer containing secondary antibodies for 2 hours at room temperature. Sections were then washed 3 x 5 min in ’washing buffer’. Nuclei were stained with Hoechst 3342 for 30 min at room temperature, followed by washing with ‘washing buffer’ (3 x 5 min) and cover-slipped with Fluoro-Gel Mounting Medium with Tris Buffer. A detailed description of the immunohistochemistry protocol is provided in the supplementary methods section.

### 4.7. Imaging and image analysis for mouse samples

Fluorescently labeled cochlear sections and whole mounts were imaged on a Leica Stellaris 5 confocal system using a 40x (1.25 NA) objective, 1X digital zoom, and z-axis spacing of 0.36 µm. To allow quantitative comparison between experimental and control conditions, images were captured using identical settings, including constant laser intensity/gain settings throughout the experiment. Cochlear mid-modiolar sections were imaged, and data from two adjacent sections were averaged, “n” refers to the number of subjects. Image analysis was performed using IMARIS (Oxford Instruments, UK) software. Cell counts and measurement of volumes were performed on maximum intensity z-projections of 3D confocal image stacks.

#### 4.7.1. Quantification of lymphangiogenesis and angiogenesis

The outline of the scala tympani (ST) at the base of the cochlea was traced, and the volume of the ST was measured. Creating a mask for individual channels, the volume of Prox1+, PECAM1+ cells was measured. The volumes of Prox1+ lymphatics and PECAM1+ blood vessels were then divided by the volumes of respective ST, expressed/10^5^ μm^3^.

#### 4.7.2. Quantification of cellular density

Images were coded with 12-digit codes containing letters and numbers; personnel involved in cell counting and volumetric analyses were blinded to the experimental conditions. The outlines of the ST, Rosenthal canal, and lateral wall at the base of the cochlea were traced, and the volumes of each area were measured. CD11c+ dendritic cells were counted. Density of Hoechst 3342+ nuclei, CX3CR1+ mononuclear phagocytes (MNP), CX3CR1-CD45+ non-MNP immune cells, CX3CR1-CD45+ CD3e+ T lymphocytes, and CX3CR1+ MHCII+ antigen-presenting cells were expressed /10^5^μm^3^. The density of CD11c+ dendritic cells, intralymphatic CD11c+ dendritic cells, was expressed/10^8^ μm^3^. Co-localized nuclei with the reporters or immunolabeled markers were used as a criterion for the identification of individual cell types.

### 4.8. Identification of lymphatics in cadaveric samples of implanted human cochlea

#### 4.8.1. Collection and preparation of human temporal bones

Temporal bone sections from an 81-year-old male with progressive bilateral sensorineural hearing loss beginning in his 50s were selected for immunofluorescence analysis. He underwent right cochlear implantation via cochleostomy with full insertion of a straight electrode (Advanced Bionics HiRes Advantage 1J. Advanced Bionics, Valencia CA) one year prior to death. The temporal bones were obtained at 12 hours postmortem, processed by standard histologic methods,(62) and sectioned in a horizontal angle at 20 um thickness. Both the implanted and contralateral, unimplanted ears were used for immunofluorescence.

#### 4.8.2 Immunohistochemistry and immunofluorescence microscopy of human temporal bones

Sections were mounted on Superfrost Plus slides, baked at 40°C overnight, and decelloidinized using saturated sodium methoxide (1:2 in methanol) for 5 minutes.(63) Sections were blocked with 5% normal horse serum and incubated overnight at room temperature with rabbit anti-LYVE1 (1:100; Abcam) and mouse anti-CD34 (1:100; Dako Omnis). Secondary antibodies goat anti-rabbit IgG (1:200; Jackson ImmunoResearch) and donkey anti-mouse IgG Alexa Fluor 568 (1:200; Thermo Fisher Scientific) were applied for 1 hour. Signal amplification used streptavidin Alexa Fluor 647 and rabbit anti-goat Alexa Fluor 488 (1:200; Thermo Fisher Scientific) for an additional hour. Nuclei were counterstained with Hoechst 33342 (1:1500; Thermo Fisher Scientific) for 15 minutes. Slides were cover slipped with anti-fade medium and imaged on a Leica TCS SP8 confocal microscope (Leica Microsystems).

### 4.9. Statistical analysis

Statistical analyses for cell count and lymphatic, vascular, and fibrotic tissue volume within the cochlea were performed using Graphpad Prism 10.4.1. Respective figure legends demonstrate the specific comparisons made. The Shapiro-Wilk or D’Agostino-Pearson test was used to determine the normality of the data. Paired t-test was used to compare the implanted and contralateral control groups. The results of the statistical analyses are described in the text. Statistical annotations (e.g., p-value) are included in the text and shown in graphs. Statistical significance was defined as p<0.05.

## Study approval

All animal experiments performed in this study were approved by the University of Iowa Institutional Animal Care and Use Committee (IACUC), according to the Guide for the Care and Use of Laboratory Animals from the Institute for Laboratory Animal Research, National Research Council, and reported following ARRIVE guidelines.

## Data availability

All data presented in this article will be shared upon reasonable request. Supplemental Materials and Methods are provided in the Supplemental Information File.

## Author Contribution

MTR, MRH, NSB, YKB, AQ, ADC: conceived of the presented idea.

MTR, IR, SMF, BJM, BH, OT, MFH, ZZ, YLE, RDG: Carried out the experiments.

MTR, SMF, IR, NAK, JO, MH, ADC, and MRH analyzed the data.

MTR prepared the figures with support from IR, ADC, and MRH

MTR wrote the manuscript with support from MRH, NSB, YKB, AQ, ADC, IR

All authors reviewed the manuscript.

## Supporting information

Supplementary Video 1- Stria Vascularis Lymphatic-PDPN

Supplementary Video 2-ScalaTympani-PDPN

Supplementary Video 3-PEPT-RWM

Supplementary Video 4- Whole mount-Thy1-PDPN

Supplementary Video 5-Synapse Lympahtics

Supplementary Video-6- Lymphatic-Dendritic Cells

## Acknowledgments

We acknowledge Linjing Xu for her continuous support in overseeing laboratory operations and coordinating resources necessary for this work.

