## Supplementary Materials for "Establishment of a cochlear lymphatic vascular network and immune surveillance system following cochlear implantation"

**Author affiliations:**

**This PDF file includes:**

Supporting text

Figures S1 to S6

Legends for Movies S1 to S6

SI References

**Other supporting materials for this manuscript include the following:**

Movies S1 to S6

### Supporting Information Text

### Supplementary materials and methods

#### Immunohistochemistry

Under ketamine (80mg/kg) and xylazine (10mg/kg) anesthesia, transcardial perfusion of ice-cold Phosphate Buffer Solution (PBS) followed by transcardial perfusion of 4% paraformaldehyde (PFA) was performed to exsanguinate mice. Harvested cochleae were fixed overnight with 4% PFA at 4°C in the dark. Following removal of excess PFA with PBS in a rotator overnight, cochleae were decalcified in 0.1M EDTA (pH 7.5) solution on a rotator for 3 days. Decalcified cochleae were rinsed with PBS, and cryoprotection of decalcified cochlea was performed using increasing (10-30%) concentrations of sucrose solutions as we have described previously[1]. Cryopreserved cochleae were infused with O.C.T. embedding medium (Tissue-TEK), mounted to the stage of a sliding block microtome (American Optical 860) stage with O.C.T. and dry ice. 30µm or 60µm thick sections parallel to the mid-modiolar plane, placed on Fisher Superfrost slides, were stored at - 20°C until immunolabeling was performed.

For immunolabeling, slides were warmed to room temperature (~20-22°C, for 10 minutes), washed (3 x 5 min each wash) with 'washing buffer' containing TBS, 0.03% Triton X-100, and 0.1% Tween-20. This was followed by permeabilization and blocking of samples in 'blocking buffer' (0.03% Triton X-100 and 0.1% Tween-20, 1% bovine serum albumin (RPI, CAS#9048-46-8), and 0.02% sodium azide (Sigma, catalog #S2002) in TBS for 2 hours. Following blocking, sections on slides were incubated in primary antibodies mixed with 'blocking buffer'. The following antibodies used in this study (with respective clone, catalog#, vendor, dilutions) are: rat anti-mouse LYVE1 (ALY7, catalog # 14-0443-82, eBioscience™, 1:200), goat anti-mouse VEGFR3/Flt-4 (catalog # AF743, R&D Systems, Bio-Techne, 1:50), Syrian hamster anti-mouse podoplanin ((eBio8.1.1 (8.1.1)) eBioscience™, catalog # 14-5381-82, eBioscience™, 1:200), rabbit recombinant anti-LYVE1 (EPR21857, catalog # ab219556, Abcam, 1:500) for human cadaveric samples, MHC Class II ((M5/114.15.2) eBioscience™, catalog # 14-5321-82, eBioscience, 1:200), mouse anti-Tubulin β3 (TUBB3) antibody (Tuj1, catalog# MMS-435P, Biolegend, 1:1000), mouse anti human myelin basic protein (MBP) (F-6, catalog# sc-271524, Santa Cruz Biotechnology, 1:100), mouse anti-rat Rip(DSHB Cat# Rip, RRID:AB_531796, 5µg/ml), Myosin 7a(catalog# 256790, Proteus, 1:150), rabbit anti-GFP (catalog# ab290, abcam, 1:2500), Syrian hamster anti-mouse CD3e ((eBio500A2 (500A2), catalog # 14-0033-82), eBioscience™), rat anti-mouse CD45 (30-F11, catalog#NB100-77417, Novus Biologicals).

After blocking and permeabilization, sections were incubated in primary antibodies in 'blocking buffer' overnight (~16 h) at 4°C. Following primary antibody application, sections were washed (3 x 5 min) in 'washing buffer', followed by incubation in blocking buffer containing secondary antibodies (Alexa Fluor conjugates 1:400, Invitrogen) for 2 hours at room temperature. Sections were then washed 3 x 5 min in 'washing buffer'. Nuclei were stained with Hoechst 3342 (10 µg/ml in TBS, Sigma) for 30 min at room temperature, followed by washing with washing buffer (3 x 5 min) and coverslipped with Fluoro-Gel Mounting Medium with Tris Buffer (catalog #17985-10, Electron Microscopy Sciences).

Beyond the general immunolabeling protocol, several exceptions were made. The concentration of the permeabilizing agents used in washing and blocking buffers varied for different antibodies used (0.5% Triton X-100 for CD3e and CD45, and 5% Tween-20 and 5% Triton X-100 for VEGFR3). All the secondary antibodies we used were derived from goat hosts(e.g., goat anti-rat, goat anti-mouse, goat anti-hamster). VEGFR3 was an exception; the primary antibody was a goat anti-mouse. The secondary antibody for the VEGFR3 antibody was a donkey anti-goat IgG conjugated with Alexa Fluor (catalog# ab150135, Abcam). To avoid the binding of the donkey anti-goat secondary to secondaries derived from goat, we used sequential labeling. Following labeling with anti-VEGFR3 primary antibody, labeling with Alexa Fluor conjugates of donkey anti-goat secondary antibody was performed. If other primary antibodies were used with the VEGFR3 antibody, labeling with those primary antibodies was performed after labeling with the VEGFR3 antibody.

**Cochlear whole mount preparation and immunolabeling**

**Cochlear whole mount preparation**

Transcardial perfusion with ice-cold PBS followed by 4% PFA was performed under ketamine (80mg/kg) and xylazine (10mg/kg) anesthesia. After overnight fixation with 4% PFA at 4°C in the dark, excess was rinsed with PBS in a rotator overnight, and cochleae were decalcified in 0.1M EDTA (pH 7.5) solution on a rotator for 3 days. Decalcified cochleae were rinsed with PBS and used for whole-mount preparation. On a silicone elastomer-coated dissection dish, the bony capsule and spiral ligament were dissected away. The cochlear whole mount was cut into apical, middle, and basal whole-mount preparations.

**Cochlear whole mount immunolabeling**

For immunolabeling, in a 48-well plate, each cochlear turn was first submerged in PBS in a separate well. Using a pipette, PBS was carefully removed and replaced with blocking/permeabilization solution (1% Triton X-100, 1% bovine serum albumin, diluted in 10 mM PBS pH 7.4) for overnight (~16 hours) incubation. Blocking/permeabilization solution was removed, and primary antibodies were applied for overnight (~16 hours), at 4°C in the dark, on a 3D rotator. The following antibodies were used for immunolabeling of the whole mounts: Myosin7a (catalog# 256790, Proteus, 1:150), mouse anti-Tubulin β3 (TUBB3) antibody (Tuj1, catalog# MMS-435P, Biolegend, 1:1000), Syrian hamster anti-mouse podoplanin ((eBio8.1.1 (8.1.1)) eBioscience™, catalog # 14-5381-82, eBioscience™, 1:200). Following incubation with primary antibodies, whole mounts were washed with washing buffer (TBS, 0.03% Triton X-100, and 0.1% Tween-20) for 3 times ( 5 minutes each wash). Secondary antibodies were applied for 2 hours in room temperature, followed by wash with washing buffer (3 X 5 minutes). Nuclei were stained with Hoechst 3342 (10 µg/ml in TBS, Sigma) for 30 min at room temperature, followed by washing with washing buffer (3 x 5 min) and coverslipped with Fluoro-Gel Mounting Medium with Tris Buffer (catalog #17985-10, Electron Microscopy Sciences).

**Cochlear implantation in the mouse model**

Cochlear implantation was performed in the left ear of the mice, using a round window approach under 1–3% inhalational isoflurane anesthesia, as we described previously.[2] Custom 3 half-banded electrode cochlear implants (Cochlear Ltd., AUS) were used. Electrode arrays were inserted to a depth of 2.25 mm. After drilling an extended bullostomy, the implant lead wires were packed in the tympanic bulla and fixed using dental cement. Around the round window, fascia was packed to prevent the leakage of perilymph.

**Validation of protocols for immunolabeling of lymphatic markers using small intestinal samples from Prox1-GFP mice**

Small intestinal samples from Prox1-GFP mice were prepared for immunohistochemistry using the same protocol described for the cochlea, excluding the decalcification step. 30µm frozen sections perpendicular to the axis of the gut lumen were labeled with antibodies against GFP, VEGFR3, LYVE1, and PDPN. VEGFR3 labeling was performed first, followed by labeling with antibodies against GFP, LYVE1, and PDPN.

**Validation of Prox1-tdTomato: PECAM1-GFP mouse lymphatic-blood vessel dual reporter**

Lymphatics and blood vessels usually run parallel to each other in most organs; distinguishing the lymphatics from the blood vessels can be challenging.[3] CD31 (PECAM1) is a marker for blood vessels that has been used to differentiate lymphatics from blood vessels.[4] Small intestinal samples from Prox1-tdTomato: PECAM1-GFP lymphatic-blood vessel dual reporter mice were prepared for immunohistochemistry using the same protocol described for the cochlea, excluding the decalcification step. 30μm frozen sections perpendicular to the axis of the gut lumen were labeled with antibodies against GFP.

**Validation of Prox1 and PDPN as markers for cochlear lymphatics**

Single-cell RNA sequencing[5] and confocal microscopy data[6], published previously, suggests that type I SGNs express a high level of Prox1. To reexamine these findings, 30μm frozen sections parallel to the midmodiolar plane from Prox1-GFP mice cochlea were labeled with mouse anti-Tubulin β3 (TUBB3) antibody (Tuj1, catalog# MMS-435P, Biolegend, 1:1000).

Within the cochlea, other than blood vessels and neurons, distinguishing the lymphatics from Schwann cells and myelin is critical; this was done by labeling cryosections of the cochlea with RIP and myelin basic protein (MBP) as markers, respectively.

**Validation of the CD11c-DTR/GFP mouse model.**

For systemic depletion of CD11+ dendritic cells, 4 ng/g body weight DT (in PBS; Sigma D-0564) was injected into CD11c-DTR/GFP transgenic mice as previously described.[7]

**Supplementary results**

**Mouse small intestinal lymphatics express Prox1, VEGFR3, PDPN, and LYVE1**

Supplementary Figure 1 shows the co-localization of lymphatic markers (Prox1, VEGFR3+ LYVE1, and Podoplanin) in small intestinal lymphatic vasculature, validating the immunolabeling protocols. Crossing lymphatic reporter (Prox1-tdTomato) mice with blood vessel reporter (PECAM1-GFP), we have generated lymphatic-blood vessel dual reporter mice.

**Prox1-expressing small intestinal lymphatics in mice are distinct from blood vessels expressing a high level of PECAM1**

In the small intestine of Prox1-tdTomato: PECAM1-GFP mice, Prox1+ lymphatic vessels could be distinguished from PECAM1+ blood vessels (Supplementary Figure 1).

**Prox1 expression in cochlear cells other than lymphatics**

Within the spiral ganglion, a major subset of Tuj1+ spiral ganglion neurons (SGNs) express Prox1+, both in the cell soma and the processes (Supplementary Figure 2).

**PDPN is not expressed on Schwann cells and myelin**

RIP+ Schwann cells are distinct from PDPN+ cells in the cochlea (Supplementary Figure 2). Moreover, PDPN does not colocalize with MBP, suggesting that PDPN is not expressed on cochlear myelin. (Supplementary Figure 2)

**PDPN+ lymphatics are adjacent to the PECAM1+ blood vessels and are distinct throughout the cochlea**

PDPN+ lymphatic vessels are observed adjacent to the PECAM1+ blood vessels throughout the cochlea: margin of the modiolus, stria vascularis, along the margin of the perilymphatic space (ST and SV) as shown in Supplementary Figure 3.

**CX3CR1+MHCII+ APCs are located near the vascular plane throughout the cochlea**

CX3CR1+MHCII+APCs are distributed throughout the cochlea near the vascular plane: margin of the modiolus, along the margins of the perilymphatic space (RWM, margins of ST and SV), as shown in Supplementary Figure 3.

**Validation of CD11c-DTR/GFP mouse model**

**CD11c+ dendritic cell depletion with diphtheria toxin:** Systemic depletion of CD11+ dendritic cells was accomplished by a single intraperitoneal injection of 4 ng/g body weight DT (supplementary Figure 4), validating the CD11c-DTR/GFP mouse model.

**CX3CR1+MHCII+ APCs and CD11c+ dendritic cell recruitment in the implanted cochlear regions other than ST**

Increased density of CX3CR1+MHCII+ cells was not observed in the spiral ligament of the base cochlea, stria vascularis ($p = 0.2003, 0.6417, 0.7464$, respectively). A trend of increase in CD11c+ dendritic cell density in the spiral ligament ($p=0.0779$), and not in stria vascularis and spiral ganglion ($p=0.4927$, 0.2268, respectively). Statistical analyses were done using paired t-test, $n=4$ for each condition (Supplementary Figure 5).

**Non-MNP immune cells and T lymphocyte recruitment in the implanted cochlear regions other than ST.**

No recruitment of non-MNP(CX3CR1-CD45+) immune cells in spiral ligament, stria vascularis, or spiral ganglion ($p=0.8542$, 0.9913, and 0.7763, respectively) was observed in the implanted cochlea. Similarly, no increase in the density of T cells (CX3CR1-CD45+CD3e+) in the spiral ligament, stria vascularis, or spiral ganglion ($p=0.9730$, 0.8688, and 0.3598, respectively) was observed. Statistical analyses were done using paired t-test, $n=4$ for each condition (Supplementary Figure 6)

**Figures**

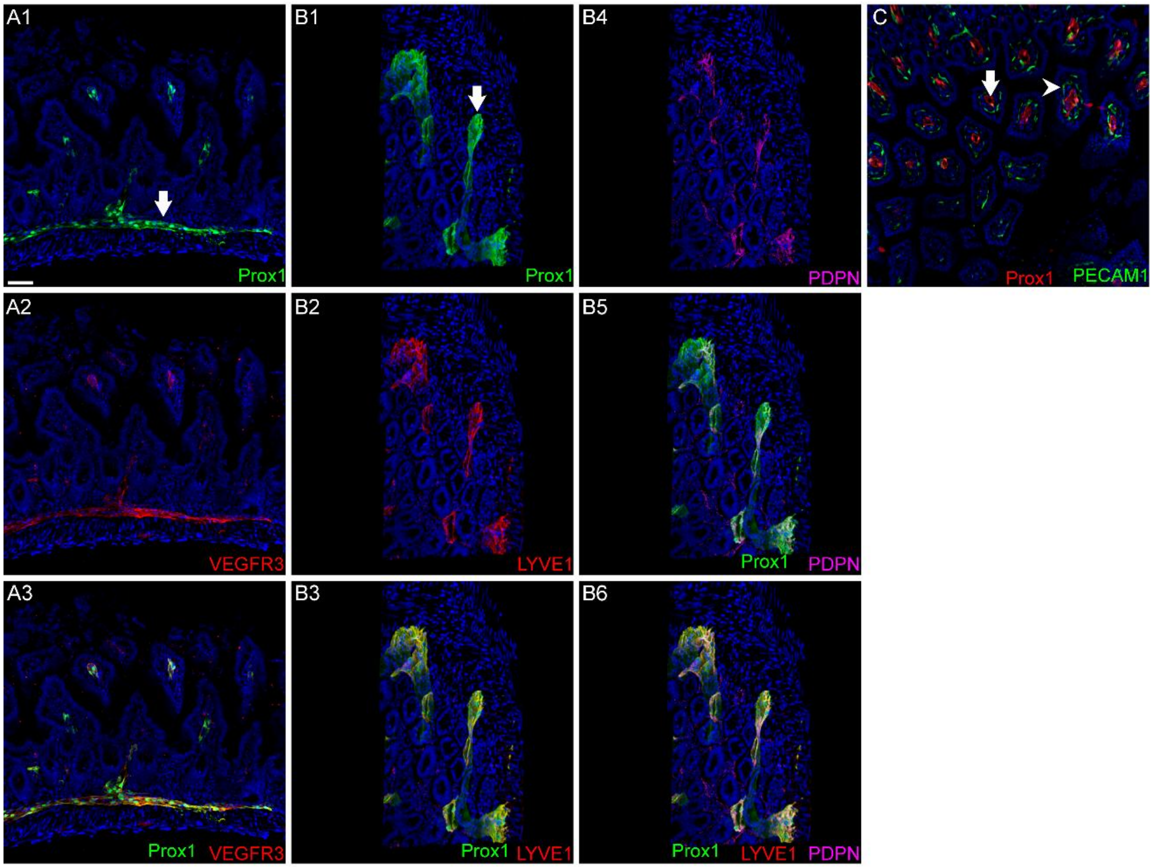


**Fig. S1. Validation of immunolabeling protocols for the lymphatic markers in the lymphatic reporter (Prox1-GFP) mice and differentiation of blood vessels and lymphatics in blood vessel-lymphatics dual reporter (Prox1-tdtomato: PECAM1-GFP) mice in the small intestine.**

**A1-A3.** 30μm-thick sections perpendicular to the long axis of the small intestine of Prox1-EGFP mice were labeled with anti-GFP and anti-VEGFR3. In the small intestine. **A1.** Prox1 expression, **A2.** VEGFR3 expression, and **A3.** Merge of **A1** and **A2** showing co-localization of Prox1 and VEGFR3 in the wall of the small intestine. **B1-B6.** Small intestinal sections of Prox1-EGFP mice were immunolabeled with antibodies against Lyve1 and PDPN. **B1.** Prox1 expression, **B2.** Lyve1 expression, and **3.** Merge of **B1** and **B2** showing co-localization of Prox1 and Lyve1 in the wall of the small intestine. **B4.** PDPN expression, **B5.** Merge of B1 and B4 showing colocalization of Prox1 and PDPN, **B6.** Merge of B1, B2, and B4 showing colocalization of Prox1, Lyve1, and PDPN. **C.** 30μm-thick sections perpendicular to the long axis of the small intestine of Prox1-tdtomato: PECAM1-GFP mice were labeled with anti-GFP antibody. Prox1+

lymphatic vessels were distinct from highly PECAM1+ blood vessels. Hoechst 3332 labeling is shown as blue in all images. Scale bar: 50μm. n=3 for each condition. Arrows indicate lymphatic vessels and arrowheads indicate blood vessels.

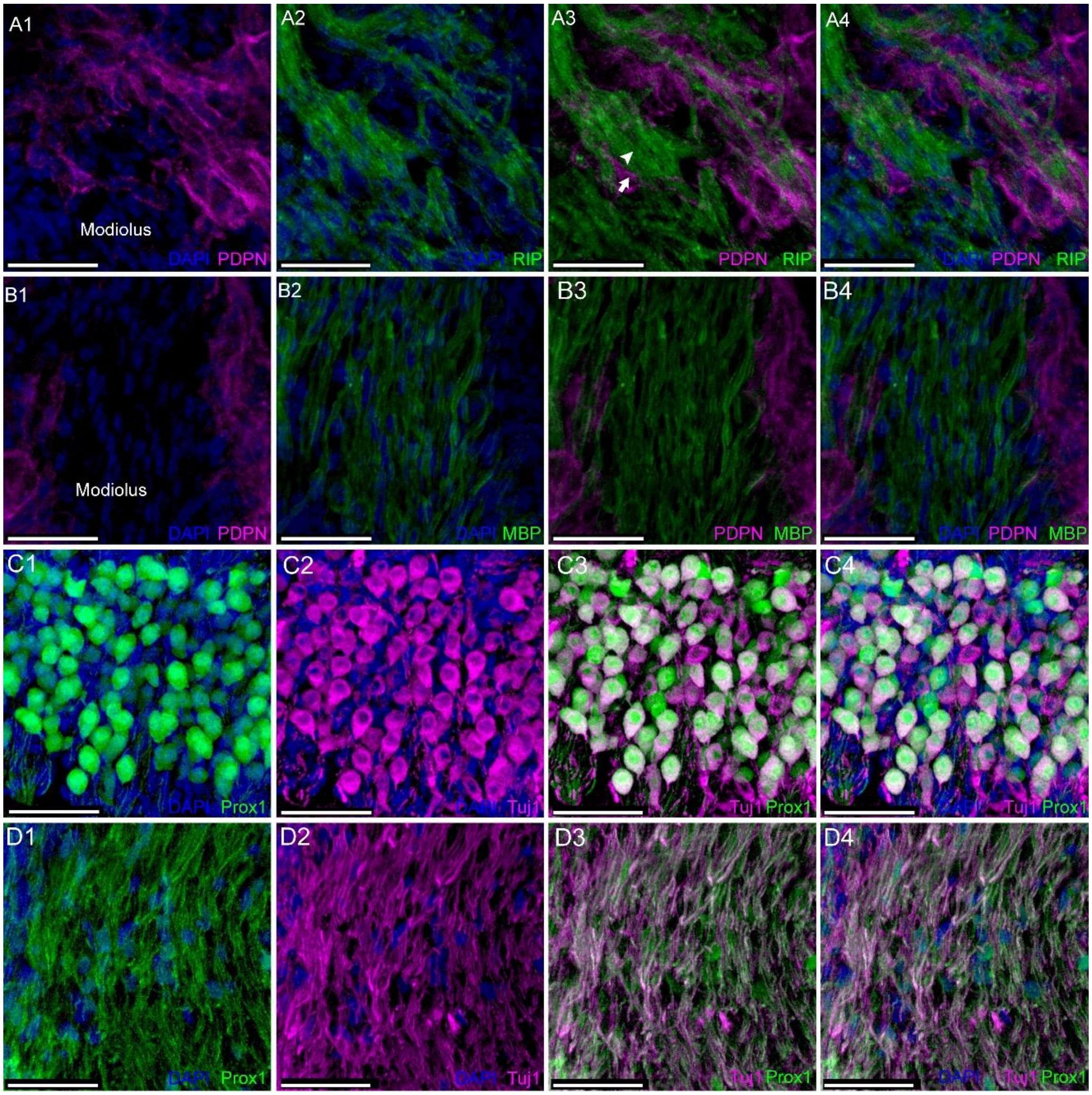


**Fig. S2. Validation of lymphatic markers in cochlear tissue. A1-A4.** To differentiate lymphatic vessels from Schwann cells, 30µm-thick cochlear sections parallel to the midmodiolar plane were labeled with anti-PDPN and anti-RIP antibodies. **A1.** PDPN, **A2.** RIP**, A3.** Merge of PDPN and RIP showing distinct PDPN+ (arrow) and RIP+ (arrowhead) cells, **A4.** Merge of PDPN, RIP, and Hoechst 3332 showing PDPN+ and RIP+ cells are different cells. **B1-B4.** To differentiate lymphatic vessels from myelin, 30µm-thick cochlear sections parallel to the midmodiolar plane were labeled with anti-PDPN and anti-MBP antibodies. **1.** PDPN, **B2.** MBP**, B3.** Merge of PDPN and MBP showing distinct PDPN+ (arrow) cells and MBP+ (arrowhead) myelin. **B4.** Merge of PDPN, MBP, and Hoechst 3332 labeling showing PDPN+ lymphatic endothelial cells and MBP+ myelin. **C1-C4** and **D1-D4.** To differentiate lymphatic vessels from spiral ganglion neurons,

30μm-thick cochlear sections parallel to the midmodiolar plane from Prox1-GFP mice were labeled with anti-Tuj1 antibody. **C1-C4.** Within the spiral ganglion, **C1.** Prox1-expressing cells within the spiral ganglion. **C2.** Tuj1 expression in SGN soma within the spiral ganglion, **C3.** Merge of **C1** and **C2**, showing that most of the Tuj1+ SGNs are Prox1+. **C4.** Merge of prox1, Tuj1, and Hoechst 3332 labeling, showing that most of the Tuj1+ SGN cells are Prox1+. **D1-D4.** within the modiolus. **D1.** Prox1-expression within the modiolus. **D2.** Tuj1 expression in the SGN neurites within the modiolus, **D3.** Merge of **1** and **D2**, showing that most of the Tuj1+ SGN neurites are Prox1+. **D4.** Merge of prox1, Tuj1, and Hoechst 3332 labeling, showing that most of the Tuj1+ SGN neurite cells are Prox1+. Scale bar: 50μm. n=3 for each condition.

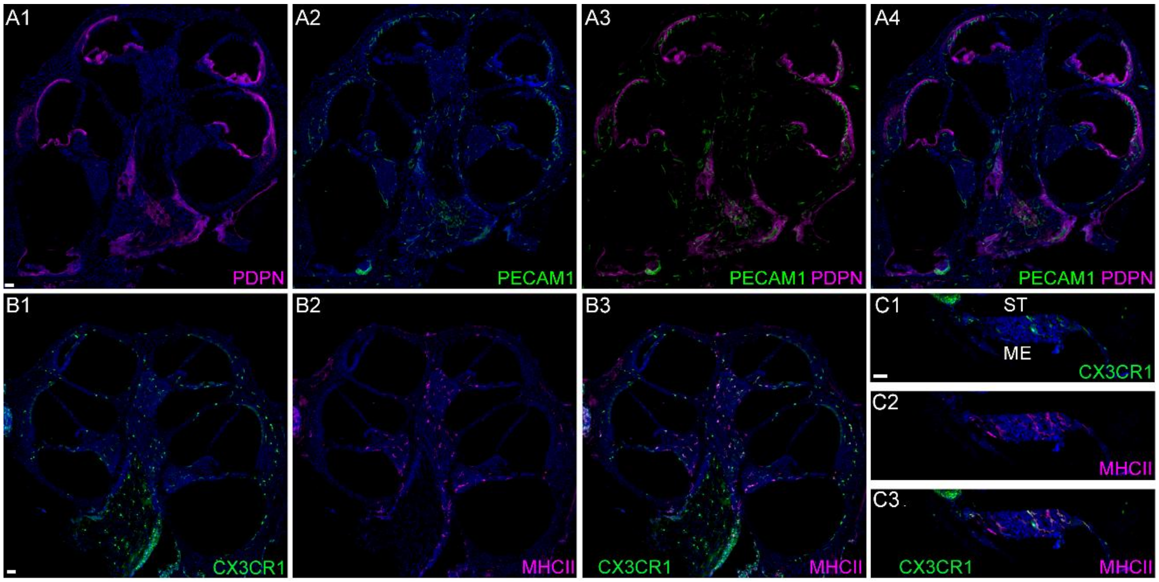


**Fig. S3. Spatial organization of lymphatics, blood vessels, and MHCII+ APCs in midmodiolar sections and round window membrane. A1-A4.** 30μm-thick midmodiolar sections of PECAM1+ cochleae were labeled with anti-PDPN antibody. Representative midmodiolar section images showing **A1.** PDPN expression throughout the cochlear tissue: spiral ganglion, outer margin of the modiolus and spiral ligament, and tissues adjacent to perilymph and endolymph. **A2.** PECAM1 expression, **A3.** Merge of **A1** and **A2** showing PDPN+ lymphatic vessels are closely related to the PECAM1+ blood vessels and are distinct. **A4.** Merge of **A1** and **A2** with Hoechst 3332 labeling showing that PDPN+ and PECAM1+ cells are distinct populations. **B1-B3.** 30μm-thick midmodiolar sections from CX3CR1$^{+/GFP}$ mice cochleae were labeled with anti-MHCII antibody. **B1.** CX3CR1-expressing cells are widely distributed in the cochlea. **B2.** MHCII expression is largely limited to the spiral ganglion, outer margin of the modiolus, spiral ligament, and tissues adjacent to the perilymph. **B3.** MHCII expression in the cochlea co-localizes with CX3CR1 expression, suggesting they are mononuclear phagocytes. **C1-C3.** 100 μm-thick sections parallel to the midmodiolar plane that include the RWM from CX3CR1$^{+/GFP}$ were labeled with anti-MHCII antibody. **C1.** CX3CR1-expressing MNPs in RWM. **C2.** MHCII-expressing APCs in RWM. **C3.** Merge of **C1** and **C2** showing colocalization of CX3CR1 and MHCII, suggesting MHCII+APCs in the round windows are MNPs. Blue indicates Hoechst 3332 stain. Scale bar: Scale bar: 50μm. n=3 for each condition.

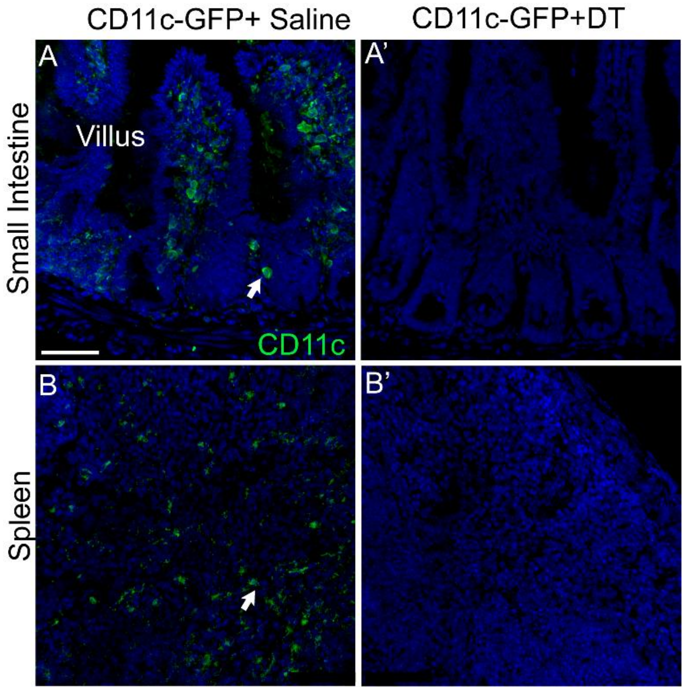


**Fig. S4. Validation of CD11c-DTR/GFP mouse model.** CD11c-DTR/GFP mice were treated with a single injection of either normal saline or diphtheria toxin (4ng/g) in normal saline. 24 hours after the injections, the ileum and spleen samples were collected. 30µm-thick sections from saline-treated **A.** Ileum and **B.** Spleen samples show CD11c+ dendritic cells (arrows), while dendritic cells are absent in DT-treated **A1.** Ileum and **B1.** Spleen. Blue indicates Hoechst 3332 stain. Scale bar: Scale bar: 50µm. n=3 for each condition.

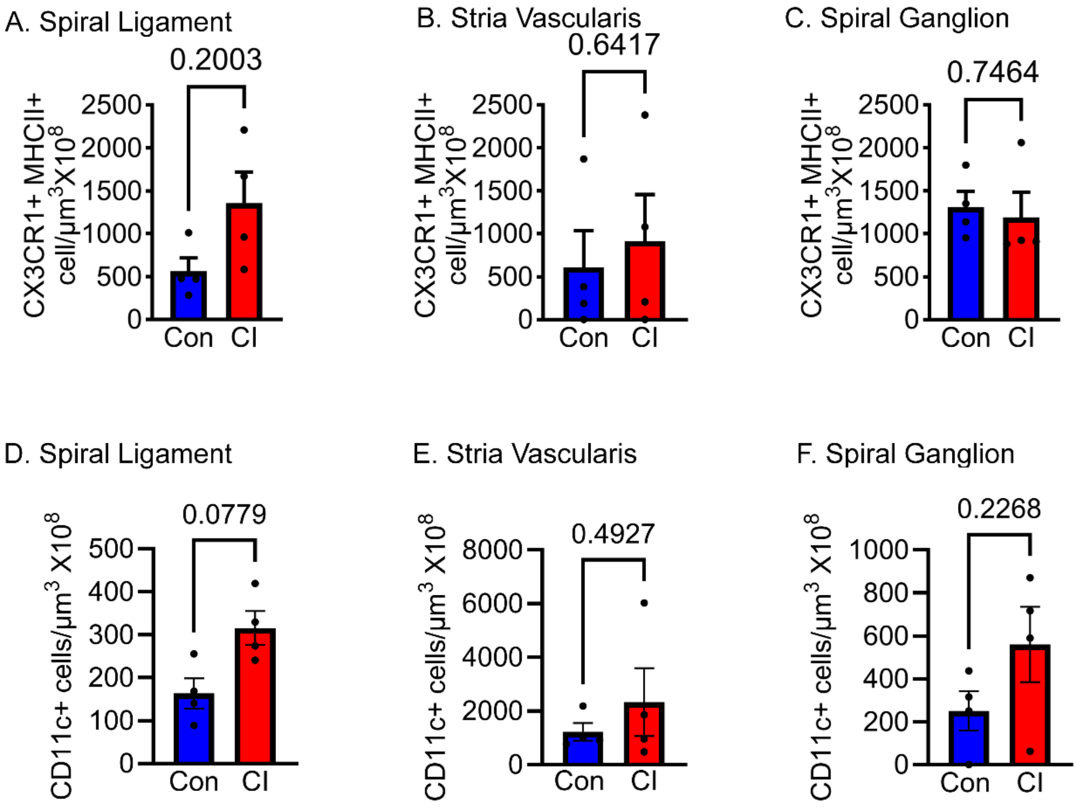


**Fig. S5. CX3CR1+MHCII+ APCs and CD11c+ dendritic cells in areas of the cochlear base other than ST.**

Twenty-eight days after cochlear implantation in CX3CR1$^{+/GFP}$ reporter mice, 30 µm-thick sections parallel to the midmodiolar plane of the cochlea were labeled with anti-MHCII antibody. Quantitative analyses of the density of CX3CR1+MHCII+ APCs (**A-C)** and CD11c dendritic cells **(D-F).** Quantitative analyses of the CX3CR1+ MHCII+ cell density in **A.** Spiral ligament, **B.** Stria vascularis, and **C.** Spiral ganglion of the base of the cochlea. Statistical analyses were performed using paired t-test. n=4 for each group. Error bar indicates SEM. p-value was shown on each graph. Quantitative analyses of the CD11c+ cell density in **D.** Spiral ligament, **E.** Stria vascularis, and **F.** Spiral ganglion. Statistical analyses were performed using paired t-test. n=3 for each group. Error bar indicates SEM. p-value was shown on each graph. Quantification of (CX3CR1-CD45+CD3e+) immune cells.

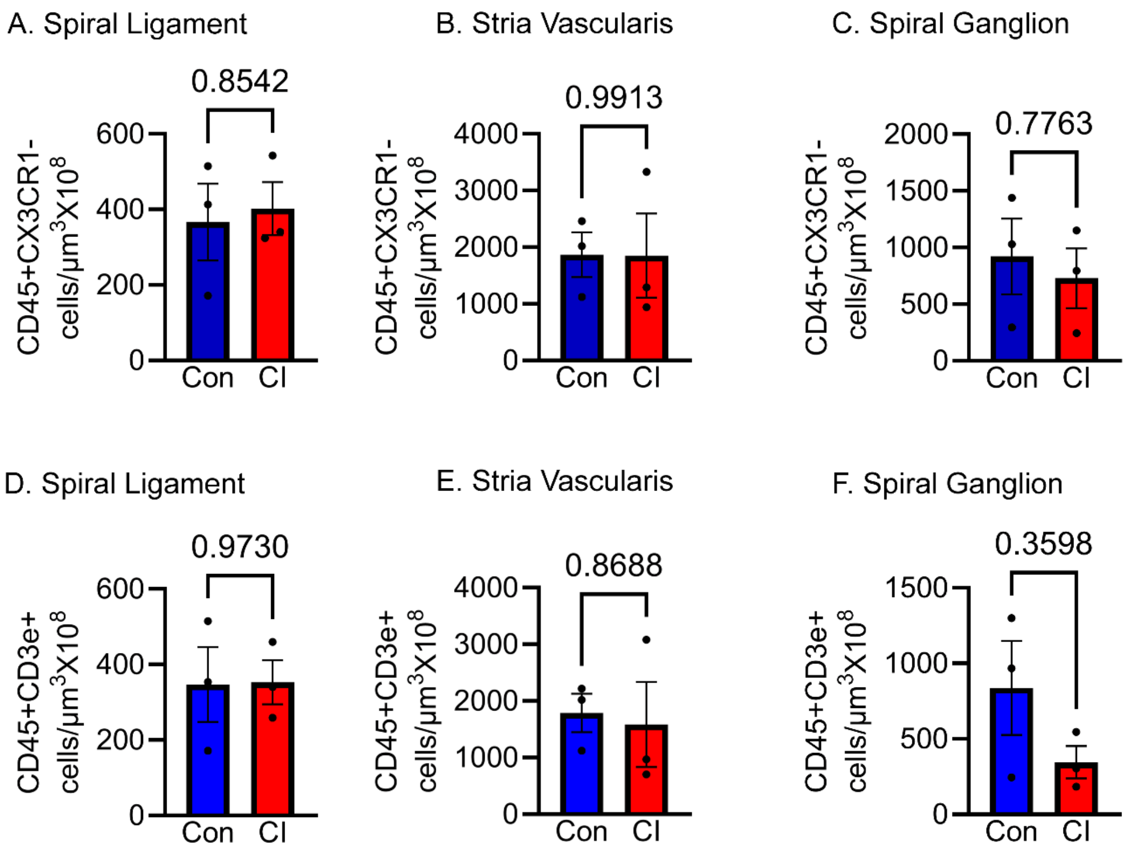


**Fig. S6. CX3CR1-CD45+ (immune cells other than MNPs) and CX3CR1-CD45+CD3e+ adaptive T lymphocytes in areas of the cochlear base other than ST**. Twenty-eight days following cochlear implantation in CX3CR1$^{+/GFP}$ reporter mice, 30µm sections parallel to the midmodiolar plane were labeled with an antibody against pan-leukocyte marker CD45 to differentiate CX3CR1+ mononuclear phagocytes and other non-mononuclear phagocytic (CX3CR1-CD45+) immune cells. Adaptive T lymphocytes were labeled with anti-CD3e antibody. Quantitative analyses of the density of CX3CR1-CD45+ (non-MNP immune cells) (**A-C)** and CX3CR1-CD45+CD3e+ adaptive T lymphocytes **(D-F).** Quantification of CX3CR1-CD45+(non-MNP) immune cells in **A.** Spiral ligament, **B.** Stria vascularis, and **C.** Spiral ganglion of the base of the cochlea. Statistical analyses were performed using paired t-test. n=4 for each group. Error bar indicates SEM. Quantification of (CX3CR1-CD45+CD3e+) adaptive T cells in **A.** Spiral ligament, **B.** Stria vascularis, and **C.** Spiral ganglion of the base of the cochlea. Statistical analyses were performed using paired t-test. n=4 for each group. Error bar indicates SEM. p-value was shown on each graph. Quantification of (CX3CR1-CD45+CD3e+) immune cells.

**Movie S1. PDPN+ lymphatics in the stria vascularis of the cochlea.** Cochlear sections (100μm) from C57B6 mice were labeled with anti-PDPN antibody. Podoplanin+ (magenta) lymphatic vessels are observed in the stria vascularis of the base of the cochlea. Hoechst 3332-labeled nuclei are shown in blue.

**Movie S2. PDPN+ lymphatics along the margin of the ST**. Cochlear sections (100μm) from C57B6 mice were labeled with anti-PDPN antibody. Podoplanin+ (magenta) lymphatic vessels are observed along the margin of the ST of the cochlea. Hoechst 3332-labeled nuclei are shown in blue.

**Movie S3. Prox1+ lymphatics and PECAM1+ blood vessels in the round window membrane.** Cochlear sections (100μm) parallel to the midmodiolar plane from lymphatic-blood vessel dual reporter (Prox1-tdttomato: PECAM1-GFP) mice were labeled with anti-GFP antibody. Prox1+ (red) lymphatic vessels are observed in the RWM that are distinct from the PECAM1+ (green) blood vessels. Hoechst 3332-labeled nuclei are shown in blue.

**Movie S4. PDPN+ lymphatics in cochlear whole mount preparation from Thy1$^{+/YFP}$ mouse**. Whole-mount cochlear preparation from Thy1$^{+/YFP}$ mouse was labeled with anti-PDPN antibody. PDPN+ lymphatic vessels (magenta) are distinct from the Thy1+ (yellow) SGNs.

**Movie S5. PDPN+ lymphatics underneath the hair cells in cochlear whole mount preparation.** Whole-mount preparations were labeled with anti-PDPN and anti-myosin 7a antibodies. PDPN+ lymphatic vessels (magenta) were observed underneath the hair cells.

**Movie S6. Intralymphatic CD11+ dendritic cells within the lumen of proliferated lymphatic vessels in the implanted cochlea**. A lymphatic-dendritic cell dual reporter (Prox1-Tdtomato: CD11c-GFP) mouse was implanted. Twenty-eight days post-CI, 30-μm cochlear sections from the implanted cochlea were labeled with an anti-GFP antibody. Prox1+ lymphatics (red) are observed around the implant track. CD11c+ dendritic cells (green) are seen inside the lumen of the Prox1+ lymphatic (red) vessels. Hoechst 3332-labeled nuclei are shown in blue.
